# Polyclonal lymphoid expansion drives paraneoplastic autoimmunity in neuroblastoma

**DOI:** 10.1101/2021.12.14.471886

**Authors:** MI Rosenberg, E Greenstein, M Buchkovich, M Mikl, A Peres, E Santoni-Rugiu, D Reshef, A Salovin, DL Gibbs, MS Irwin, A Naranjo, I Ulitsky, PA de Alarcon, V Weigman, G Yaari, JA Panzer, N Friedman, JM Maris

## Abstract

Neuroblastoma is a lethal childhood solid tumor of developing peripheral nerves. Two percent of children with neuroblastoma develop Opsoclonus Myoclonus Ataxia Syndrome (OMAS), a paraneoplastic disease characterized by cerebellar and brainstem-directed autoimmunity, but typically with outstanding cancer-related outcomes. We compared tumor transcriptomes and tumor infiltrating T- and B-cell repertoires from 38 OMAS subjects with neuroblastoma to 26 non- OMAS associated neuroblastomas. We found greater B- and T-cell infiltration in OMAS- associated tumors compared to controls, but unexpectedly showed that both were polyclonal expansions. Tertiary lymphoid structures (TLS) were enriched in OMAS-associated tumors. We identified significant enrichment of the MHC Class II allele HLA-DOB*01:01 in OMAS patients. OMAS severity scores were associated with the expression of several candidate autoimmune genes. We propose a model in which polyclonal autoreactive B lymphocytes act as antigen presenting cells and drive TLS formation, thereby crucially supporting both sustained polyclonal T-cell-mediated anti-tumor immunity and paraneoplastic OMAS neuropathology.

## Introduction

Immune surveillance, the idea that the immune system plays an important role in eliminating tumor cells, was first introduced over a hundred years ago by Paul Erlich (Erlich P 1909). The complex process of immune modulation (“immune editing”) of tumor growth is robustly supported by multiple mouse studies that demonstrate spontaneous tumor generation and metastasis (reviewed in e.g. Swann and Smyth, 2007) This editing process involving early elimination of tumor cells, an equilibrium of evolving tumor and immune restriction, and eventual tumor escape, finds abundant support in human disease as well. The complete spontaneous regression of certain types of neural crest cancers, like neuroblastoma and melanoma (McGovern, 1975) demonstrates the potential of effective immune surveillance in eliminating cancer in humans. Careful investigation of rare patient populations that exhibit particularly effective deployment of immune surveillance is therefore warranted.

In rare instances in a naturally occurring setting, individuals with solid tumors develop autoimmunity triggered by the tumor, a condition termed paraneoplastic autoimmune disease. Many of these paraneoplastic diseases involve self antigens that are expressed in endogenous tissue of the central nervous system (CNS), causing severe neurological symptoms that range from psychosis (e.g. NMDA-receptor encephalitis, driven by teratoma; reviewed in Dalmau, et al 2019) to motor deficits, mood and behavioral changes, paralysis, and other symptoms (e.g. limbic encephalitis associated with non-small cell lung cancer [NSCLC], Shen et al 2018). The autoimmunity is presumed to be driven by a shared epitope between tumor and brain (Graus et al, 2004). Consistent with an important role for immune mechanisms in controlling tumor growth, patients with paraneoplastic diseases often have better tumor-related outcomes than patients with the same cancer but no autoimmune component (Darnell, R.B. and Posner, J.B. 2003; Smith and Stehlin 1965; Byrne and Turk 2011; Nordlund,et al 1983). Improved tumor outcomes may arise in the context of complete or partial tumor elimination; paraneoplastic disease may persist even in the absence of remaining tumor cells.

A hallmark of adaptive immunity is the remarkable combinatorial potential of lymphocytes, which are able to generate diverse antigen receptors, permitting broad and potent immunity. However, the same diversity that protects from a broad array of foreign antigens carries greater risk for autoimmunity. Therefore, in mammals, negative selection in the thymus and the bone marrow is needed to cull self-reactive immune receptors to prevent targeting of self, causing autoimmunity. The paradox of paraneoplastic disease, then, is that patients with autoimmunity possess a broader repertoire of immune reactivity with which to restrict or eradicate solid tumors than patients with proper immune selection, even as it leads to pathology of native tissue. Further evidence of this tenuous relationship is the observation that cancer patients treated with checkpoint inhibitors often develop autoimmunity (Zekeridou and Lennon, 2019; Valencia-Sanchez and Zekeridou 2021). Understanding how the delicate balance between powerful anti-tumor immunity and deleterious anti-self pathology is achieved is of critical importance in improving immunotherapy strategies for treatment of a wide range of cancers. The molecular analysis of anti-tumor immunity in rare patients with paraneoplastic disease is therefore of great interest.

Both antigen reactivity in neuroimmunity (e.g., NMDA receptor encephalitis; Sansing et al 2007) and immune repertoires in solid tumors (e.g., metastatic breast cancer; De Mattos-Arruda et al 2019) have been separately investigated. But to date, to our knowledge, no study has linked molecular characterization of tumor and its immune infiltrate with the paraneoplastic autoimmune phenotypes of the same patients, to permit elucidation of the immune process underlying paraneoplastic disease. Integrated analysis of paraneoplastic disease-associated tumors offers a unique setting for the evaluation of systemic immune features driving both powerful anti-tumor immunity and often severe native tissue pathology.

Pediatric Opsoclonus Myoclonus Ataxia Syndrome (OMAS) is a rare but devastating autoimmune disorder characterized by sudden onset of uncontrollable, irregular, multivectorial eye movements, myoclonic jerking of the limbs, ataxia, and disordered mood/behavior in a previously well child (Kinsbourne 1962). These prominent neurological symptoms, which often result in lifelong sequelae, often precipitate diagnosis of the underlying tumor. OMAS is often associated with neuroblastoma, a solid tumor of the peripheral sympathetic nervous system that arises from the neural crest during development, but can also occur when no tumor is detectable. Most OMAS patients have localized, low-risk neuroblastoma disease, infrequent *MYCN* amplification (a strong negative prognostic determinant for neuroblastoma associated with low MHC expression; Bernards et al 1986), and often harbor other genomic copy number profiles that ordinarily accompany higher risk tumors, but that are nevertheless favorably resolved (Hero et al 2018). Importantly, as with other paraneoplastic diseases, patients with OMAS and neuroblastoma have better tumor outcomes than even low-risk neuroblastoma patients without OMAS (Altman and Baehner 1976). Here, we carried out a systematic study of OMAS-associated neuroblastoma tumors accrued on prospective Children’s Oncology Group (COG) clinical trial ANBL00P3 (de Alarcon et al 2018) to define the mechanisms for improved anti-tumor immunity as well as molecular correlates of the neuroimmune disease phenotype in neuroblastoma patients with OMAS.

## Results

### Tumor gene expression profiling shows highly diverse tumor lymphoid infiltrate

To identify gene expression differences underlying differential anti-tumor immunity and neuroreactivity, we performed RNA sequencing on the 38 archival primary neuroblastoma samples from patients with OMAS treated on COG clinical trial ANBL00P3 (de Alarcon et al 2018), with 13 low-risk and 13 high-risk (7 with *MYCN* amplification) neuroblastomas from age-matched patients without OMAS, obtained through the COG neuroblastoma biology study ANBL00B1 as comparators. RNA quality was poor in many of these archival samples, necessitating use of an exome capture RNA sequencing protocol for this study (Schuierer et al 2017). Differential expression analysis was consistent with significant lymphoid infiltrate in the OMAS tumors, as expected, but showed enrichment of memory B and T cells, and not antibody secreting plasma cells as we expected (**Figure 1A-B, Table S1**). Among the most differentially expressed genes between OMAS neuroblastomas and low-risk non-OMAS neuroblastomas were *CD22* and *BANK1*, both of which modulate B cell activity, and *CCRL1*, a regulator of immune and cancer cell migration. Notably, OMAS-associated neuroblastoma showed significant increased differential expression of *TCF7*, a marker of stem-cell like CD8+ T cells and regulator of autoimmunity (**Table S1,** reviewed in Escobar et al 2020). In contrast, *GLUD2*, which has been reported to be an OMAS autoantigen (Berridge et al 2018), was not significantly differentially expressed in our OMAS-associated neuroblastoma dataset (**Figure 1A**). Highly expressed outlier genes in OMAS compared to non-OMAS also included *CR2*, a complement receptor that is expressed on dendritic cells and on B cells where it enhances binding of B cells to immune complexes and BCR signaling in autoimmunity (Kulik et al 2019). In line with prominent B cell infiltration, we observed significant differential expression of B cell chemokine, CXCL13, and its receptor, CXCR5 (**Figure 1A**). Gene set enrichment analysis using ENRICHR (Chen et al 2013) showed several hallmarks of T cell activation and differentiation, Th17 subtype specification, and B cell activation among functions of genes upregulated in OMAS tumors (**Figure 1C**). GSEA of genes significantly less expressed in OMAS compared to non-OMAS neuroblastoma highlighted extracellular matrix, including low expression of *NCAN*, a CNS-specific matrix protein, and synthesis and metabolism of chondroitin sulfate and dermatan sulfate, two extracellular matrix proteins important for neural crest cell migration and reported to have immune modulatory properties (**Figure 1D**; Su et al 2017).

**Figure 1.**
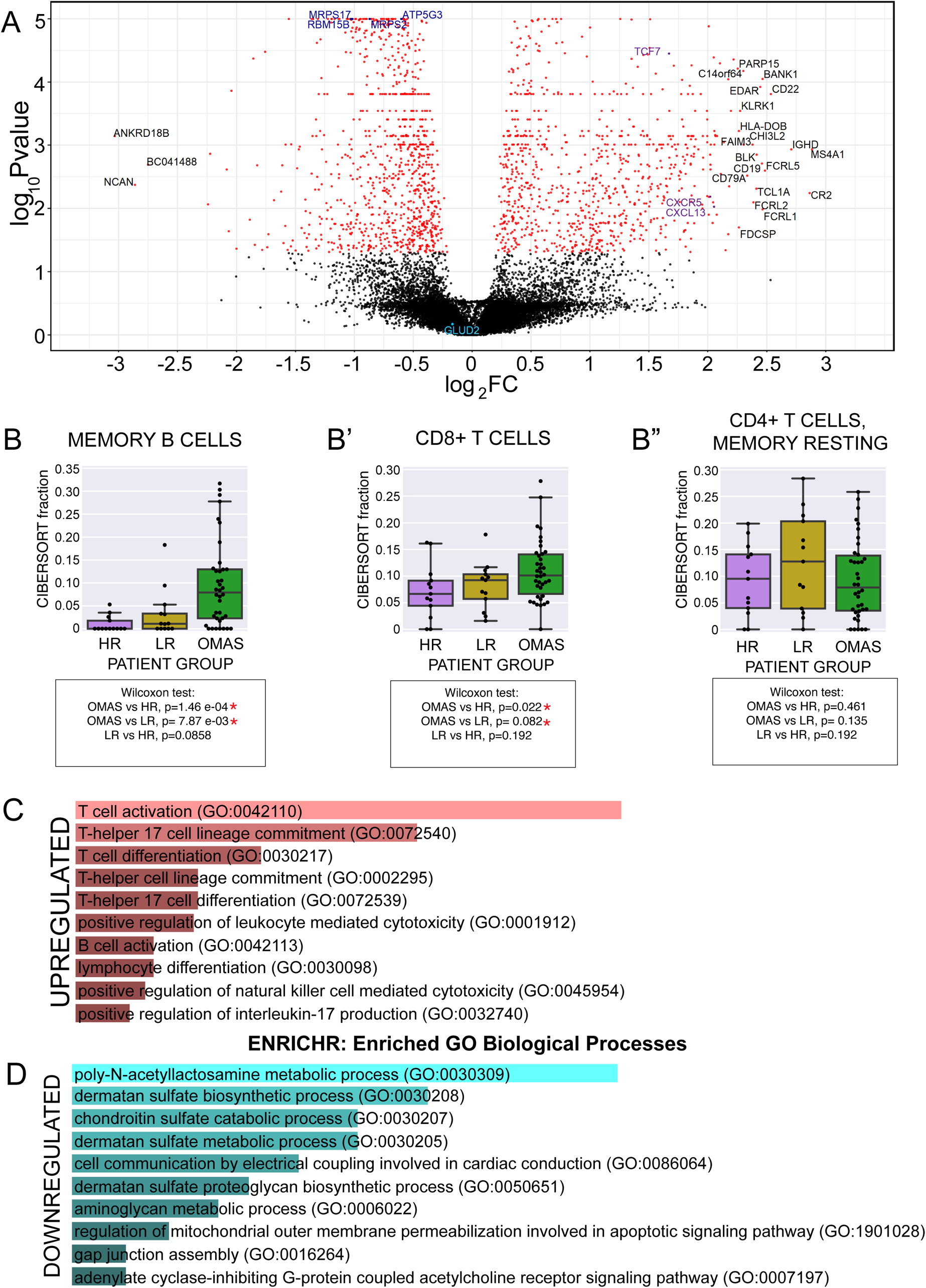
RNA-seq analysis highlights enhanced lymphocytic infiltration and activation in OMAS-associated neuroblastoma compared to control neuroblastoma. (**A**) A volcano plot comparing expression (Log2 fold change) of transcripts (as dots) in OMAS-associated neuroblastoma compared to non-OMAS neuroblastoma. X axis indicates enrichment of expression in OMAS. Significance of differential expression (LogP value) is given on the Y axis. Adjusted P value <0.05 indicated in red. Gene names in black are given for genes with expression differences of greater than Log_2_(2.25 fold) between groups. Gene names referred to in subsequent analyses labeled in light and dark blue and purple. **B-B”**) CIBERSORT analysis of gene expression values from tumor RNA-seq permit estimates of immune cell fractions in OMAS vs control neuroblastoma tumor infiltrate, including (B) memory B cells; (B’) CD8+ T cells; and (B”) Resting CD4+ T cell fractions. **(C-D**) ENRICHR analysis of significantly differentially expressed genes. Genes with *≥*2 fold difference in expression between groups were used as input for gene set enrichment analysis using ENRICHR. GO terms for Biological processes for over-represented in upregulated genes (C; red) and downregulated genes (D; blue) are shown, as bar graphs of combined significance score.

**Table 1.** Enrichment of HLA alleles in OMAS compared to control neuroblastoma patient groups. Two different models were used to test for enrichment of HLA alleles that may contribute to OMAS autoimmunity. **(A)** Allele model. This model assesses occurrence of each HLA allele in the pool of total alleles found in patients of one subtype compared to another subtype. Allele frequency calculated as # of observed alleles/total number of alleles in that population pool (2x # samples). **(B)** Population model. This model for enrichment tests for each HLA allele in patients from each population compared to another. Here, the number of patients containing the allele, regardless of copy number, is compared to the total number of patients in the pool. The total sample size for each population= the number patients; patients homozygous for the allele are counted only once.

| ALLELE<br>MODEL |  |  |  |  |  |  |  |  |
| --- | --- | --- | --- | --- | --- | --- | --- | --- |
| HLA<br>Allele | OMAS<br>freq | LR<br>freq | HR<br>freq | NonOMS<br>freq | OMASvLR<br>FDR | OMASvHR<br>FDR | OMASvnon-OMAS FDR | LRvHR<br>FDR |
| L*01:01 | 0.67 | 0.43 | 0.27 | 0.28 | 0.38 | 8.21E-08 | 5.90E-08 | 1 |
| L*01:02 | 0.28 | 0.09 | 0.09 | 0.00 | 0.49 | 0.0056 | 0.0016 | 1 |
| DOB*01:01 | 0.63 | 0.26 | 0.38 | 0.36 | 0.01 | 0.0118 | 0.0016 | 1 |
| DRB1*13:02 | 0.12 | 0.04 | 0.04 | 0.03 | 1 | 0.28 | 0.16 | 1 |
| C*04:01 | 0.21 | 0.11 | 0.09 | 0.09 | 1 | 0.19 | 0.16 | 1 |
| DRB1*01:01 | 0.2 | 0.09 | 0.09 | 0.09 | 1 | 0.28 | 0.18 | 1 |
| POPULATION<br>MODEL |  |  |  |  |  |  |  |  |
| HLA<br>Allele | OMAS<br>freq | LR<br>freq | HR<br>freq | NonOMS<br>freq | OMASvLR<br>FDR | OMASvHR<br>FDR | OMASvnon-OMAS FDR | LRvHR<br>FDR |
| L*01:01 | 0.89 | 0.44 | 0.35 | 0.35 | 0.04 | 1.79E-07 | 5.69E-08 | 1 |
| L*01:02 | 0.50 | 0.11 | 0.16 | 0.16 | 0.09 | 0.0064 | 0.0015 | 1 |
| DOB*01:01 | 0.71 | 0.33 | 0.41 | 0.40 | 0.21 | 0.10 | 0.04 | 1 |
| DRB1*01:01 | 0.39 | 0.19 | 0.16 | 0.16 | 1 | 0.14 | 0.07 | 1 |
| DRB1*13:02 | 0.24 | 0.07 | 0.07 | 0.06 | 1 | 0.18 | 0.10 | 1 |
| DQB1*05:01 | 0.50 | 0.33 | 0.25 | 0.25 | 1 | 0.18 | 0.10 | 1 |
| DQB1*06:04 | 0.16 | 0.04 | 0.03 | 0.03 | 1 | 0.18 | 0.10 | 1 |
| DQA1*01:01 | 0.42 | 0.22 | 0.20 | 0.19 | 1 | 0.22 | 0.13 | 1 |
| DQA1*05:01 | 0.42 | 0.26 | 0.21 | 0.21 | 1 | 0.22 | 0.15 | 1 |

### OMAS associated tumor gene expression reveals increased inflammation

We next explored immune landscape signatures derived from OMAS transcriptomes compared to non-OMAS tumor samples (**Figure 2A**). OMAS neuroblastomas showed significantly higher mean expression of CD8, B cell score, cytotoxic lymphocyte immune signature (CLIS), T cell co-stimulatory molecules, CD28, markers of activation, (CTLA4), and exhaustion, (PD1). These data are consistent with previously published reports of increased lymphocytic infiltration in OMAS tumors (Fukushima et al 2017; Gambini et al 2003; Cooper et al 2001), and also, with the enhanced T cell activation we show here by transcriptome profiling. OMAS samples segregate into roughly three subgroups: one with higher expression of immune gene features, one with more moderate expression, and one in which OMAS samples cluster together with high risk non-OMAS neuroblastoma samples exhibiting low immune marker scores (**Figure 2A**). The lone high-risk, *MYCN* amplified OMAS-associated neuroblastoma in the present cohort, PARSCY, did not appear in this cluster.

**Figure 2.**
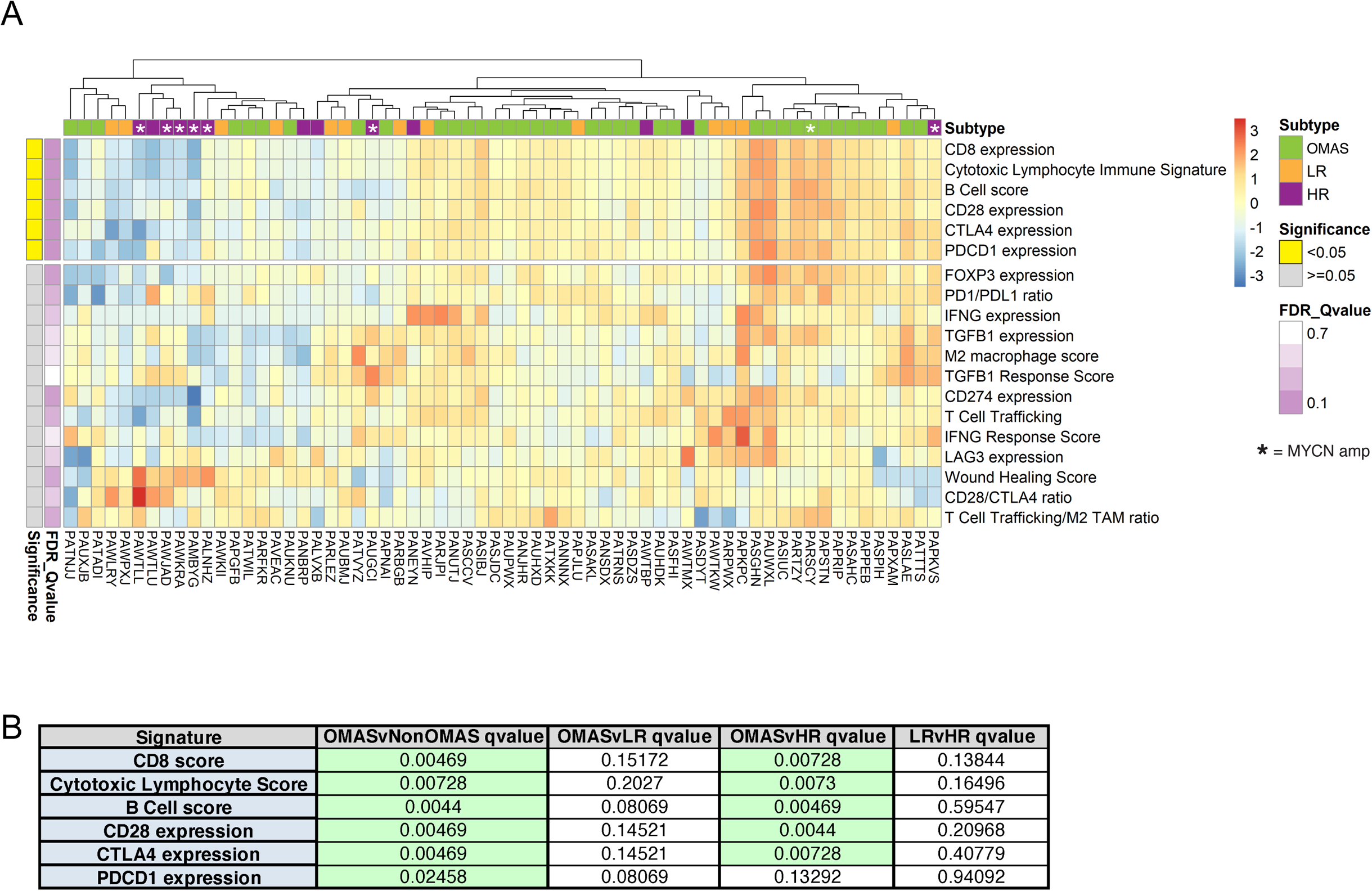
Immune landscape signature defines distinctive immune features of OMAS-associated neuroblastoma. Immune signatures were detected in each sample from RNA-seq gene expression as previously described (Jones et al 2020). Immune signature scores are weighted averages of (log) expression levels of genes within each signature. **(A)** Patients are ordered by immune score relative to mean centered values within this cohort, and clustered according to similarity of scores across signatures. Immune subtypes were tested for enrichment among the OMAS population compared to non-OMAS neuroblastoma, using a Fisher’s exact test with correction for multiple testing using Benjamini-Hochberg. Immune features that are statistically different between OMAS and non-OMAS are plotted in the upper box. Patient subtype is indicated by color at the top (green-OMAS, orange-LR non-OMAS, purple-HR non-OMAS), patient sample ID indicated below along the X axis. “*” indicates NMYC-amplified tumor. **(B)** Adjusted p values (q values) from statistical tests of enrichment for immune signature features in OMAS and plotted in the upper box of panel A are indicated. Significant values are shaded bright yellow at left margin.

Finally, to probe more deeply the differences in the tumor microenvironment in OMAS patient samples, we adapted a recently developed approach for classification of immune responses to tumor using RNA expression (Thorsson et al 2018; **Figure 3A**). We found that dominant immune signaling pathways in OMAS tumors were significantly different from either low-risk or high-risk neuroblastomas. Fifty percent of OMAS-associated neuroblastoma were classified as “IFNγ- dominant”, a classifier phenotype that predicts association with strong CD8+ signal and greatest TCR diversity, while only 10-15% of non-OMAS tumors had this classification (**Figure 3B,C**). Indeed, we observed an increased fraction of CD8^+^ T cells in OMAS tumors, as estimated from RNA-seq data using CIBERSORT (**Figure 1B**) (Chen et al, 2019; Newman et al 2015). The C2 classifier phenotype and the IFNγ and CLIS features of the immune landscape signature (**Figure 2**) converge on the strong differential T cell signature in OMAS associated tumors. In contrast, 50% of HR neuroblastomas were classified as “wound-healing dominant”, a phenotype associated with a high proliferative index and angiogenic gene expression, as well as Th2 cell bias, and importantly, a poor overall prognosis (Thorsson et al, 2018; **Figure 3B,C**). We observed a small but significant increase in the proportion of OMAS tumors over non-OMAS tumors classified as C3, or “inflammatory” subtype, a classification associated with lower levels of cell proliferation, aneuploidy and somatic copy number variation, and superior outcomes (sCNV; OMAS vs non-OMAS, FDRq =0.046).

**Figure 3.**
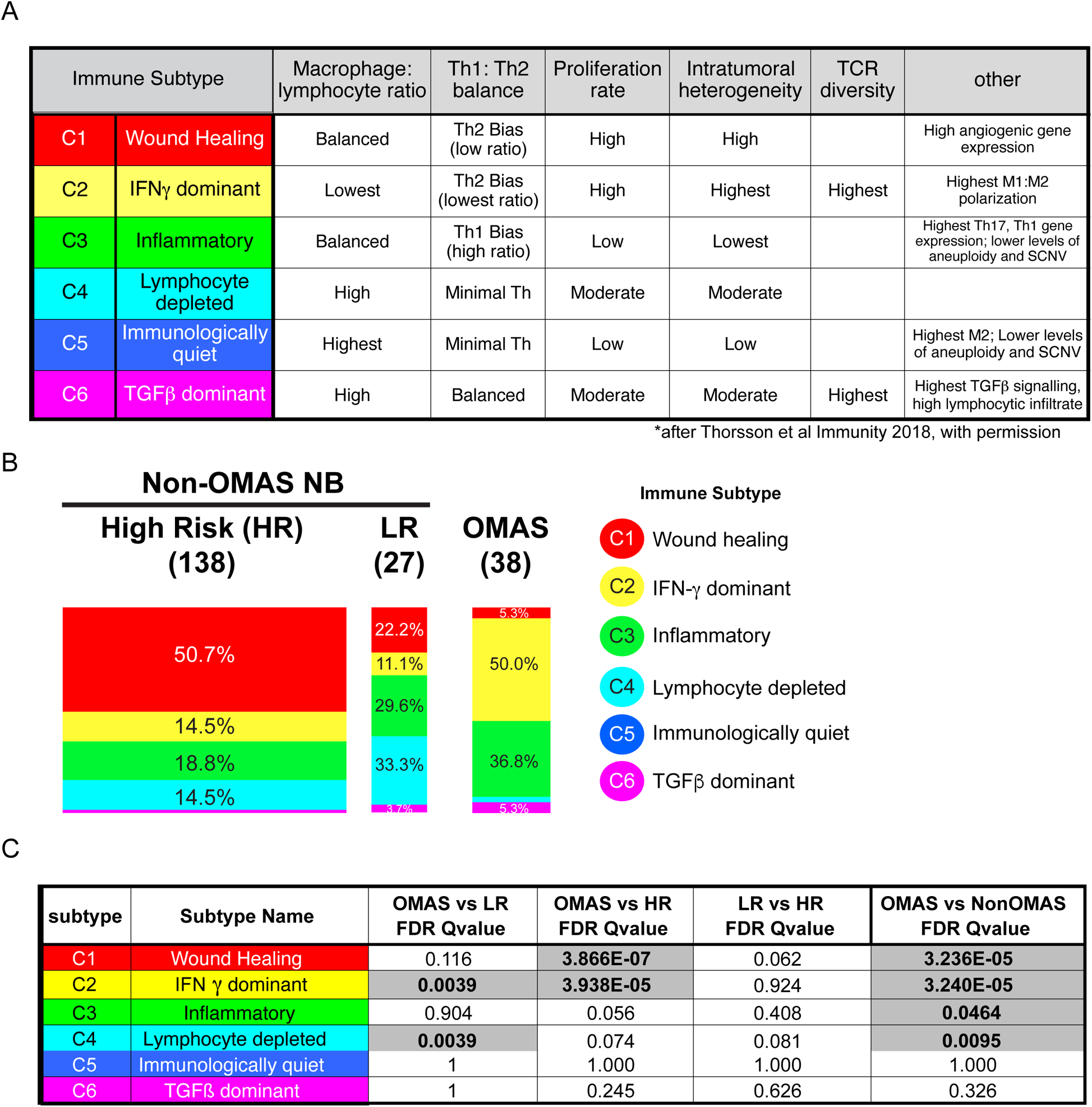
Cancer immune subtype classification identifies dominant immune signaling pathways in neuroblastoma patient cohort. Immune subtype classifications were applied using normalized RNA-seq (log) expression levels for each patient, as previously described (Thorsson et al 2018). **(A)** Features of immune subtypes. Distinctive features of immune response correlated with each subtype C1-C6 based on meta-analysis of TCGA cancer dataset are indicated. **(B)** Distribution of immune subtypes in OMAS and control neuroblastomas in this cohort. **(C)** Enrichment of immune subtypes in OMAS relative to other control neuroblastoma patient groups are indicated. Significant values are shaded in grey.

Global gene expression profiling and clustering of OMAS vs non-OMAS neuroblastomas appear to be driven by the degree and type of their immune infiltrate. Therefore, we also used a machine learning classifier, XGBoost (Chen and Guestrin 2016), to determine whether a distinguishing gene expression profile of OMAS-associated neuroblastoma could be identified. The algorithm was able to clearly distinguish OMAS from non-OMAS (auROC=0.94; **Figure S1A,D**), and to distinguish OMAS from either high risk (auROC=0.69; **Figure S1B,E**) or low risk neuroblastoma (auROC=0.69; **Figure S1C,F**) to a lesser degree. It is noteworthy that the classification was driven by very few genes, as opposed to a broader gene expression signature. The top 10 features that were, on average, most important for the correct prediction of patient population (**Figure S1G**) included *MRPS2*, *RMB15B*, and *MRPS17*, encoding mitochondrial proteins. Lower expression of each of these genes drives the prediction towards OMAS (**Figure 1A**, dark blue; **Figure S1A,G**), which may be attributable to an increased proportion of dead or dying cells in the OMAS samples (Ilicic et al 2016).

Though OMAS-associated neuroblastoma is expected to have a modest mutational load typical of neuroblastoma (Pugh et al 2013; Brady et al 2020), we investigated potential sources of neoepitope variation that could contribute to increased anti-tumor immunity by analysis of SNV burden using RNA sequencing data. We identified 94 genes enriched for SNV variation that were significantly different between OMAS and non-OMAS samples, of which 47 genes are significant compared to HR alone (FDR q value <0.20; **Table S2**). However, we did not identify any single source of epitope variation in all patients that obviously underlies the observed immune response to OMAS-associated neuroblastomas.

### Expression of several CNS cell surface genes are correlated with OMAS disease severity

OMAS can present with neurological symptoms ranging from mild to severe and debilitating, and a semi-quantitative grading system has been devised (De Grandis et al 2009). We examined whether gene expression or immune features in the tumor correlated with disease severity scores of OMAS collected at the time of diagnosis. Expression of two neuronal cell surface receptors: the serotonin receptor, *HTR6*, and an alpha 2 adrenergic receptor, *ADRA2C,* correlated significantly with severity of OMAS neuroimmune symptoms (**Figure S2**). The gene *NCAN*, a CNS specific extracellular matrix protein whose expression has been linked to malignant behavior of neuroblastoma (Su et al 2017), also correlated significantly with OMAS neuroimmune symptoms. This candidate is also noteworthy, since it is among the most differentially expressed genes in OMAS tumors compared to non-OMAS tumors (**Figure 1**). Expression of additional genes relevant for adhesion of neurons and leukocytes (*DSCAML1, MADCAM1*) was also significantly correlated with OMAS symptom severity.

### MHC Class II alleles distribution in OMAS-associated neuroblastoma

Susceptibility to many autoimmune diseases has been linked to genes encoded by the major histocompatibility complex (MHC) (reviewed in (Dendrou et al 2018). We inferred HLA types from tumor derived RNA using the *RNA Access* library platform, as described above. We then used HLAprofiler, a published computational tool for HLA calling from RNA-seq data with >99% concordance with direct DNA sequencing (Buchkovich et al 2017). To establish background HLA allele frequencies in neuroblastoma, we inferred HLA types from a large set of neuroblastoma transcriptomes from the NCI-TARGET neuroblastoma dataset (Pugh et al 2013) using HLAprofiler and compared allele frequencies from our OMAS cohort to non-OMAS controls from this study and non-redundant set of TARGET transcriptomes. The non-classical class II allele, HLA-DOB*01:01, was significantly enriched in OMAS (FDR q Value=0.002; **Table 1 and Table S2**). HLA-DO regulates MHC class II peptide loading and is almost exclusively expressed in B cells and in thymic epithelial medullary cells but not other professional APCs (Karlsson et al 1991). HLA-DOB was also significantly differentially expressed in OMAS tumors compared to non-OMAS tumors (**Table S1**). Use of a less stringent FDR threshold of 0.2 to allow for discovery of additional alleles from our relatively small cohort of cases (with false discovery rate of 20%) allowed detection of HLA-DRB1*01:01 as being enriched in our OMAS cohort, consistent with a previous report (Hero et al 2005; HLA DRB1*01; FDR q=0.18), as well as HLA-DRB*13:02 (FDR q= 0.16) and one MHC Class I allele, HLA-C*04:01 (FDR q=0.16). The most skewed HLA alleles we identified were two different alleles of the MHC Class I pseudogene HLA-L. HLA-L is highly expressed in EBV transformed B-cells, however its functional significance is unknown.

### Tumor infiltrating T cells exhibit greater antigen receptor diversity in OMAS-associated neuroblastoma

A link between the OMAS autoimmune response and improved anti-tumor immunity would predict that the repertoires of tumor infiltrating T cells and B cells would be strongly shaped by OMAS causative antigen(s). We hypothesized that the OMAS tumor lymphocytic infiltrate would be predominantly oligoclonal. We used genomic DNA from tumors to sequence TCR β and the immunoglobulin heavy chain (IgH) repertoires (Robins et al 2009; Carlson et al 2013), and analyzed lymphocyte repertoires from 31 OMAS samples, and 13 LR and 13 HR control samples. We analyzed in-frame sequences corresponding to the TCRβ and IgH CDR3 regions, which provide most of the antigen binding specificity to the receptor, and therefore are used as a proxy for antigen specificity of each receptor type in this analysis. OMAS-associated neuroblastoma TCR repertoires were significantly larger than those recovered from HR neuroblastoma samples (**Figure 4A**, FDR q=0.001), and 2-fold larger than low-risk neuroblastoma samples (**Figure 4A**, FDR q= 0.071). These T cell number estimates based on genomic DNA sequencing of TCRβ repertoires are consistent with RNA-seq estimates of higher T cell numbers in OMAS samples, using differential marker gene expression and as detected by CIBERSORT (**Figure 1**), and the immune landscape signature (**Figure 2**).

**Figure 4.**
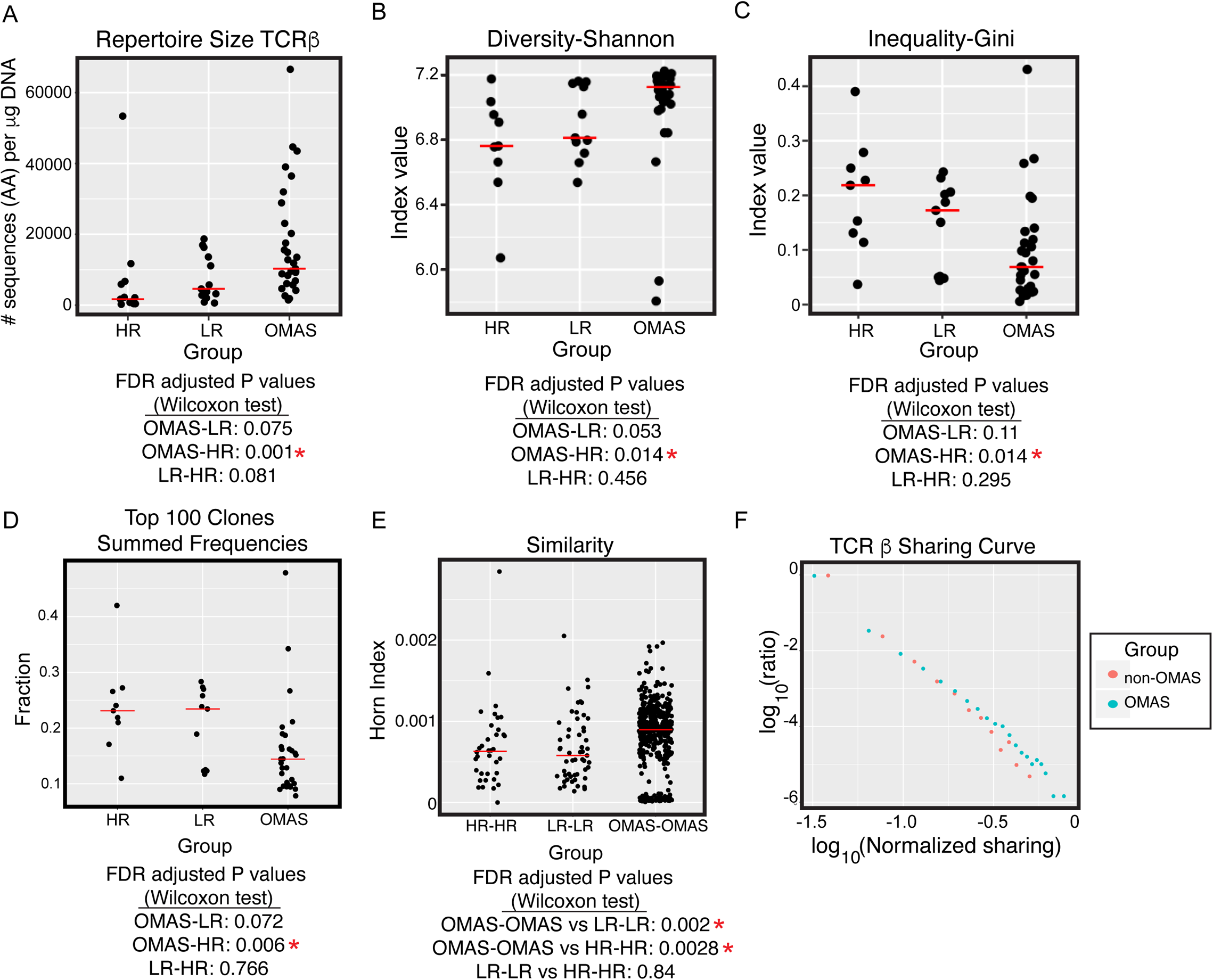
OMAS tumor infiltrating T cell receptor repertoire analysis reveals significant diversity and small clones, with limited similarity and sharing of primarily public sequences. (**A**) Shannon diversity index of OMAS-associated and non-OMAS-associated neuroblastoma TCRβ repertoires. Repertoires were subsampled to 1382 sequences and Shannon index computed. Average over 100 iterations plotted for each patient. Median value indicated in red. (**B**) Gini index of evenness of OMAS-associated and non-OMAS neuroblastoma TCRβ repertoires. Average over 100 iterations plotted for each patient. Median value indicated in red. (**C**) Sums of clonal frequencies for top 100 clones of TCRβ repertoires. Cumulative individual frequencies of top 100 clones in each patient repertoire were summed and plotted as a single point. Median value in each patient subgroup indicated in Red. (**D**) Horn index. Scatter plot of Horn Index values (after downsampling with replacement to 1382 sequences) of pairwise comparisons between patients within each patient subgroup. Average result over 100 iterations is plotted. Median value indicated in red. I Sequence sharing in OMAS and non-OMAS patient TCRβ repertoires. Sharing value computed for each CDR3 as two numbers: 1) sharing level between OMAS samples (the number of OMAS samples that have the given CDR3) and sharing level in the control samples (sum of sharing levels within the HR and LR samples). Each dot represents the fraction of sequences in the given sharing level, normalized by the number of samples in each group. The figure is in log10-log10 scale.

We next evaluated the diversity and clonality of the TCRβ repertoires. To minimize the effect of sample size on diversity estimates, we down-sampled all repertoires to a common size (reducing the analysis to 49 samples out of 57 total). We then computed Shannon entropy (a measure for diversity) and Gini index (a measure for clonal inequality) for each sample, averaging over 100 iterations of subsampling. We found that OMAS repertoires are significantly more diverse than either high-risk (**Figure 4B**; FDRq=0.014) or low-risk (FDRq=0.053), while the latter non-OMAS cohorts were similarly diverse (p= 0.456). The higher diversity of OMAS TILs is in line with the observation of increased TCR diversity for tumors of immune classifier subtype C2 (Thorsson et al 2018), which is dominant among our OMAS samples (**Figure 3C, D**).

TCR repertoires within OMAS samples had significantly lower Gini indices, a measure of clonal evenness, than non-OMAS neuroblastoma samples (**Figure 4C**), indicating more even distribution of clone sizes, without considerable expansion. In accordance with their Gini indices, we found that the summed frequencies of the top clones were also significantly lower in OMAS compared to either low risk or high risk (**Figure 4D, Figure S3**). Together, these results invalidated our original prediction of oligoclonality in TIL repertoires and instead support the notion that OMAS-associated neuroblastomas harbor diverse, polyclonal repertoires of T cells.

### TCRβ repertoires from OMAS patients share highly public TCR CDR3β sequences

We then compared similarity of tumor infiltrating TCR repertoires from patients with and without OMAS using the Morisita-Horn index to capture the degree of similarity between samples. To minimize the bias of the larger repertoire size of OMAS samples, the Horn index was calculated after down-sampling repertoires to a common size (1382 sequences, which reduced the total cohort to 49 total samples). Figure 4E shows average Horn Index values for pairwise comparisons between patients in each class; greater index value indicates greater similarity. OMAS repertoires exhibited greater similarity than control neuroblastoma repertoires, though the Horn index values are relatively small, suggesting that the sharing is limited. To rule out more subtle, convergent specificity, we conducted an independent and somewhat more permissive search for similarity across repertoires. We used TCRdist (Dash et al 2017), an algorithm that scores occurrence of a TCR in different repertoires within a specified distance threshold of permitted substitutions or gaps, with concomitant scoring penalties, and assesses overlap of clusters of similar TCRs with a specified cohort. TCRdist also did not return any significant similarity of shared, cohort-specific sequences (**Table S3,** sheets 1-3).

Plotting the histogram of sharing for the two groups nevertheless supports a somewhat greater similarity between OMAS samples (**Figure 4F**). We captured the difference between the two sharing distributions by comparing the number of sequences that appear in a single repertoire (“private”) to the number of sequences shared by at least two samples. Out of 691,960 unique amino acid sequences in OMAS samples, 4.9% of them were shared by two or more OMAS patients. In contrast, out of 208,357 unique sequences in non-OMAS neuroblastoma controls, only 3.2% were shared by two or more patients (Fisher test; p<2.2×10^-16^). Greater sharing among OMAS patients is also evident from the sharing distribution with the OMAS distribution uniformly above the control sharing distribution (**Figure 4F**). Together, these observations suggest that the observed greater sharing between OMAS repertoires is likely driven by a small number of TCR sequences.

It is noteworthy that most of the highly shared CDR3β sequences in OMAS repertoires, as well as in non-OMAS neuroblastoma repertoires, are also highly shared in PBMCs of healthy donors (found in >75% of 786 repertoires reported in Emerson et al, 2017) suggesting that these are likely public sequences (**Figure 4E, Table S3** sheet 4, “OMAS highly shared Public”). A subset of OMAS-associated shared TCRs that are less shared among non-OMAS neuroblastoma patients in our cohort (“OMAS overshared”) and another subset that are enriched in non-OMAS neuroblastoma (“Control overshared”) are summarized in **Table S3**. While their specificity may still be unknown, some shared enriched TCRs in different patient subgroups have been previously reported in other disease contexts, which may yield additional insights from the literature.

### Diversity of B cell IgH repertoire is associated with improved OMAS tumor-related outcomes

B cell infiltration of solid cancers generally has positive prognostic value, and yet the role of B cell infiltration of solid tumors is far less well understood than that of CD8+ T cells (reviewed in Nelson 2010). In contrast, the central role for B cells in OMAS neuropathology is underscored by the efficacy of the anti-CD20 antibody rituximab in mitigating neurological symptoms in OMAS (Pranzatelli et al 2006; Wilbur et al 2019). Given the significant B cell infiltrate evident from tumor RNAseq, we predicted an oligoclonal response which would be evident in analysis of IgH repertoires from OMAS associated tumors.

As with TCRs, OMAS-associated neuroblastomas had larger BCR repertoires than either HR (p= 0.01) or LR (p=0.12); non-OMAS neuroblastoma repertoire sizes were not significantly different in size (HR-LR: p=0.46) (**Figure 5A**). As for TCRβ, we calculated the Shannon diversity index for all IgH repertoires after down-sampling to a common size. We found that OMAS BCR repertoires were significantly more diverse than in control neuroblastomas (**Figure 5B**). Shared clinical features of OMAS may be associated with dominance of a few large clones responding to the OMAS antigen(s) in the CNS compartment, which we predicted would also be represented in OMAS tumors. We therefore investigated the clonal structure of OMAS tumor repertoires. LR and HR tumors both possessed larger clones than patients with OMAS (**Figure 5C**; OMAS-HR, FDRq=0.011; OMAS-LR, FDRq=0.14; LR-HR, FDRq= 0.16). We also examined whether VH or JH differed in gene or gene family usage or in CDR3 length in OMAS. However, only very low frequency events were detected as significant (**Figure S4**).

**Figure 5.**
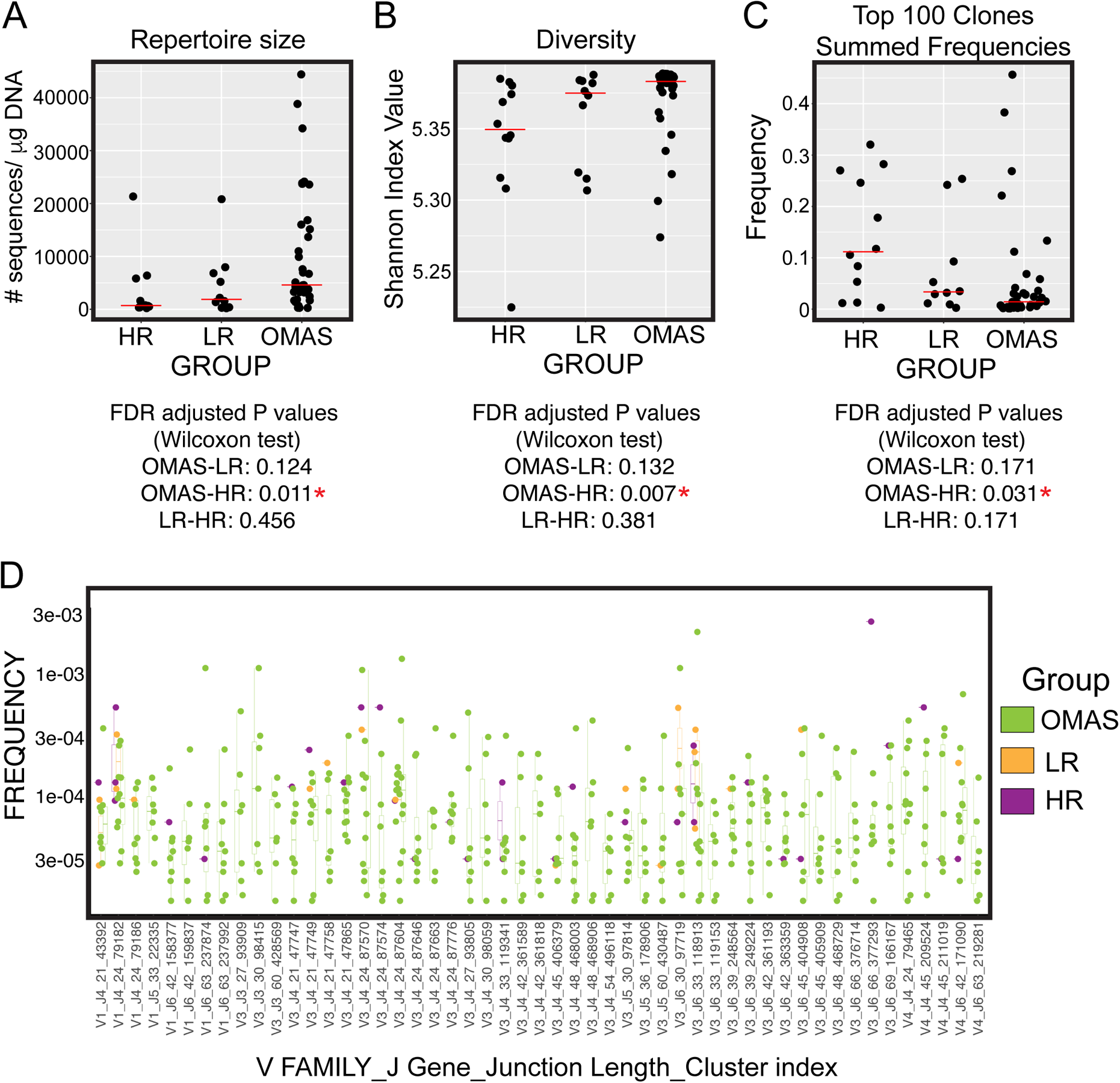
IgH repertoire analysis of tumor infiltrating lymphocytes reveals greater diversity, reduced clonality of OMAS-associated neuroblastoma BCR repertoires. (**A**) Shannon diversity index of OMAS and non-OMAS-associated neuroblastoma IgH repertoires. Mean index value after 100 iterations of downsampling and index calculation is plotted as one point for each patient. Red line indicates median for each patient group. (**B**) Clone size of OMAS and non-OMAS-associated neuroblastoma TIL repertoires. Summed frequency of top 100 clones in each patient is given as a point. Red line indicates median value for each patient group. (**C**) IgH clusters enriched in OMAS. Clusters of IgH sequences with at least 85% sequence similarity, and comprising at least 7 OMAS patients and not more than 2 LR or HR patients are shown, with V family, J family junction length and cluster index indicated.

### OMAS enriched clones exhibit similar sequence features

Owing to the uneven sizes of the OMAS and control repertoires, and to the small repertoire sizes for all samples, we were unable to test whether clones observed only in OMAS repertoires are truly OMAS-specific. Figure 5D highlights clusters of sequences possessing 85% sequence similarity and shared by at least 7 OMAS patients, grouped by VH and JH gene usage and junction length. Several sequences were not observed at all in HR patients in this study; many were also only shared by a single LR patient. We also characterized numbers of somatic mutations in IgH V genes, as a marker of somatic hypermutation in B cell clones. Increased numbers of mutations would be acquired in mature germinal center B cells and are used as a proxy for B cell clonal selection. We detected a few significant increases in somatic mutation frequency in the IGHV genes in OMAS compared to LR or HR (**Figure S4F**, stars). However, we cannot infer any biological relevance of these mutation rates from the current cohort.

Taken together, the significantly greater B cell infiltration in OMAS tumors was characterized by paucity of large clonal expansions. The B cell infiltrates were significantly more polyclonal in OMAS compared to control neuroblastoma patients. This diversity, as well as our limited number of control samples and their small repertoire sizes precluded nomination of any specific BCR clone or sequence as specifically correlated with OMAS or anti-tumor immunity.

### OMAS tumors contain germinal centers and exhibit apparent neuronal localization of tumor infiltrating lymphocytes

Histological examination revealed numerous tertiary lymphoid structures (TLSs) resembling germinal centers (GCs) in 10 of 14 OMAS tumors available for evaluation (**Figure 6A; Figure S5**) usually accompanied by widespread interstitial lymphocyte infiltration. In contrast, 2 of 6 non-OMAS low-risk neuroblastoma and 1 of 5 non-OMAS high risk neuroblastoma displayed similar structures. The TLSs contained dense cores of CD20+ B cells surrounded by CD3+ T cells, and were easily distinguished from neighboring tissue by morphology using differential interference contrast (DIC) or bright field microscopy. Using an antibody against Ki67, a marker of cell proliferation, we observed relatively few Ki67-positive cells within putative GCs in OMAS tumors (**Figure S5B**). We also noted localization of B cells and T cells to putative neuronal processes within small patches of differentiating neuroblasts in OMAS tumors (**Figure 6B, Figure S5C**). This often included B cells at the center with T cells enriched nearby.

**Figure 6.**
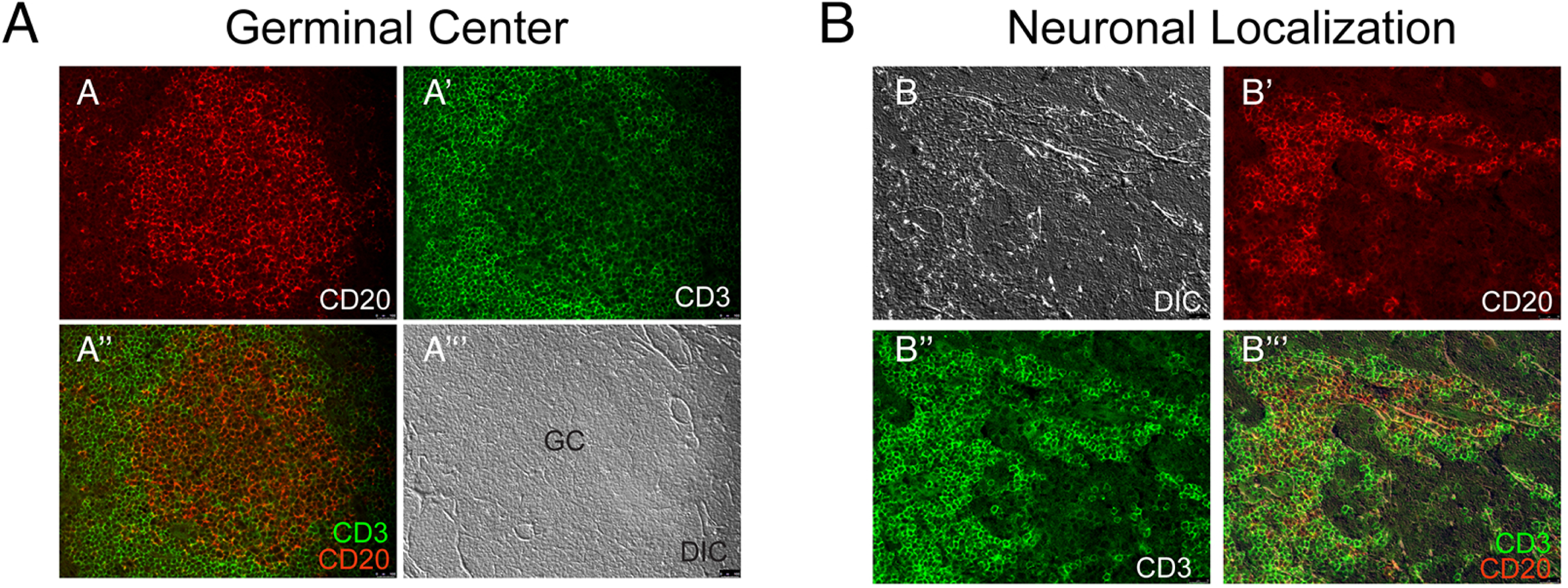
Lymphocyte localization to tertiary lymphoid structures resembling germinal centers, and to cells resembling ganglia, in OMAS-associated neuroblastoma masses. (**A-A”’**) Representative tertiary lymphoid structure in OMAS-associated neuroblastoma, containing (**A**) B cells (anti-CD20+; red), (**A’**) T cells (anti-CD3+; green). (**A”**) Merge of green and red channels; (**A”’**) DIC image of the same field. (**B-B”’**)

## Discussion

Here, we sought to understand the underlying mechanisms of neuroblastoma-associated autoimmunity with a characterization of tumors from patients enrolled on the only prospective OMAS clinical trial reported to date (de Alarcon et al, 2018). To our surprise, we found that the robust immune cell infiltrate is dominated by polyclonal B and T cells, absent the identification of a unifying single antigenic stimulus, as has been seen in other paraneoplastic diseases (e.g., NMDAR encephalitis [Dalmau J et al, 2007; Jones BE et al 2019). We confirmed a major role for autoreactive B cells in neuroblastoma associated OMAS, and here highlights a major role for T cells in antitumor reactivity and likely neuropathology, importantly, in the context of TLSs. We also identify an MHC Class II allele, HLA DOB*01:01, as significantly enriched in OMAS compared to NB controls.

In this work, we compared OMAS to non-OMAS neuroblastoma, with additional contrast of OMAS vs low-risk neuroblastoma, to highlight the influence of paraneoplastic autoimmunity on superior anti-tumor reactivity, and to pinpoint foci of OMAS neuroimmune targeting. While no clear, single molecular target of neuroimmunity emerged, we identified four conspicuous differences between OMAS and non-OMAS-associated neuroblastomas, also remarkable in OMAS vs low-risk NB, which align with reported signatures from solid tumor literature as having positive prognostic value. These same features accompany tissue infiltrates in human autoimmune disease, supporting their relevance for CNS tissue pathology in OMAS, and supporting their centrality in a systemic OMAS disease process. These are: 1) increased numbers and activation of B cells in tumor infiltrate, rich in memory B cells, 2) localization of B cell infiltrate to tertiary lymphoid structures rich in T cells, 3) polyclonality of lymphocytic infiltrate and 4) differential expression of *TCF7*, CXCR5 and CXCL13. These features accompany significant TCR and BCR diversity in OMAS tumors compared to controls, which is a defining feature of OMAS associated NB, but one whose relevance to disease outcomes is less clear. Combining these observed differences with insights from both cancer and autoimmunity, we propose a framework to explain how systemic autoimmunity drives superior tumor outcomes and neurological damage in OMAS.

For anti-tumor immunity, it is striking that the same defining features of OMAS mirror the immune characteristics of tumors from other cancers with positive response to immune checkpoint blockade, including another neural crest-derived cancer, melanoma (Helmink et al 2020; Cabrita et al 2020). While CD8+ T cells are considered the workhorses of tumor destruction, OMAS tumors exhibit greatest differences in B cell numbers, exceeding even low-risk neuroblastomas which also have excellent outcomes; OMAS tumor transcriptional profiles suggest enrichment of memory B cells, and histopathological evaluation finds that OMAS neuroblastomas contain more tertiary lymphoid structures (Table S7; Fukushima et al 2017; Gambini et al 2003). The presence of TLSs in tumors has been identified as a strongly predictive prognostic factor for positive tumor outcomes across cancer types (Ruffin et al 2021, Dieu-Nosjean et al 2016), and has been noted after successful cancer immunotherapies (reviewed in Trub and Zippelius 2021; Sautes-Fridman et al 2019). While it is not known what drives TLS formation, we observe differential expression of B cell chemokine CXCL13 and its receptor CXCR5 in OMAS tumors compared to non-OMAS (**Table S1**), two features correlated strongly with ectopic lymphoid structure formation in a variety of settings in both cancer and autoimmunity (reviewed in Kazanietz et al 2019). A *TCF7+* T cell subset has independently been identified as enriched in TLSs of an oral solid tumor, and predictive of positive tumor outcomes (Peng et al 2021). Consistent with both TLS enrichment and superior outcomes in OMAS associated tumors, *TCF7* is strongly differentially expressed in OMAS tumors. The signature of TCF7, CXCR5/CXCL13 in B cell rich TLSs was found as a predictor of survival in melanoma independently of all other variables (Cabrita et al 2020), underscoring their importance for outstanding tumor outcomes across cancer types. The extreme diversity and significant polyclonality of OMAS lymphocytic infiltrate is not easily nor universally aligned with solid tumor outcomes in other cancers, where diversity and clonality may accompany either positive or negative outcomes. For example, increased diversity of TCR repertoires has been linked to improved tumor outcomes after immune checkpoint blockade (Robert et al 2014; Valpione et al, 2021), but increased, not diminished, clonality was predictive of positive outcomes (Valpione et al 2021). Taken together, we therefore identify TLS with diverse polyclonal lymphocytic infiltrate, and strong expression of B cell chemokines and *TCF7*, as the signature of paraneoplastic autoimmunity most prominently associated with superior tumor outcomes in OMAS.

These same features of OMAS-associated neuroblastoma have also been noted in pathological tissue infiltrates in human autoimmune disease (reviewed in Jones and Jones, 2015). B cells and their trafficking to sites of inflammatory cytotoxicity are emerging as central to disease severity in autoimmunity, as well. In autoimmune encephalitis caused by multiple sclerosis (MS), B cell follicles and B cell chemokine CXCL13 expression are enriched at brain lesions associated with severe, progressive disease presentation (Magliozzi et al 2007), while loss of CXCL13 in a mouse MS model mitigates severe disease phenotypes (Bagaeva et al 2006). Similarly, high levels of CXCL13 have been found in inflamed synovia of patients with severe rheumatoid arthritis (RA; Bugatti et al 2014), while loss of CXCR5 in mouse models of RA reduced joint damage and impaired TLS formation (Wengner et al 2007, Moschovakis et al 2017). In previous studies of OMAS, high levels of CXCR5 and CXCL13 were noted in cerebrospinal fluid (CSF) of patients with OMAS, correlated with increased disease severity (Pranzatelli et al 2012). This chemokine/receptor pair mediates migration of B cells, which we now link to trafficking both to tumor and CNS in OMAS. The presence of tertiary lymphoid structures accompanies disease severity and target tissue damage in a range of autoimmune diseases (reviewed in Pipi et al 2018), and predict similar pathology in the CNS of OMAS patients, though TLSs in the brains of living OMAS patients cannot be investigated.

Tertiary lymphoid follicles are sites of antigen presentation that arise in peripheral tissues upon chronic inflammatory stimulation that often accompanies autoimmunity or infection (reviewed in Trub et al 2021, Sautes-Fridman et al, 2019). They support memory B cell formation, auto-reactive T- and B- cell activation, and can also lead to production of high affinity antibodies, via plasma cell differentiation. Germinal centers are those TLSs with mature plasmablasts that have undergone somatic hypermutation to produce high affinity, presumably cytotoxic antibodies (Shlomchik and Weisel, 2012). In OMAS associated neuroblastomas, we identify TLSs and memory B cell enrichment, as well as B cell follicles rich in T cells. However, Ki67, a histological marker of proliferation often associated with clonal expansions of antibody rearranged B cells, was largely absent from these structures in our cohort (**Figure 6; Figure S5**). Furthermore, in our data, we observe an absence of dominant species of expanded B cell clones in IgH repertoire analysis, and the absence of strong BLIMP1 expression, a marker of germinal centers, alongside strong differential expression of CD22, a B cell marker that is not expressed in mature plasma cells. Together, these findings could suggest either that we have observed a snapshot of TLS maturation that precedes a complete germinal center reaction, or that the antibody function of OMAS B cells may not be its essential one.

We propose that the critical function of B cells in OMAS tumor and CNS immunity is not only the production of pathogenic antibodies but as potent antigen presenting cells (APCs) in long-lived tertiary lymphoid structures. In the context of neuroimmunity, B cells function crucially as APCs in a lupus-prone mouse model (Giles et al., 2015) and in the EAE murine model of multiple sclerosis (Molnarfi et al 2013). EAE model mice expressing the MOG-specific B cell receptor but unable to secrete antibodies are fully susceptible to EAE induction by MOG in an MHC Class II dependent manner (Molnarfi et al 2013). Since antigen-experienced B cells of animals with autoimmunity function as APCs, and may spontaneously drive TLS formation, these interactions result in CNS targeting and T cell mediated cytotoxicity in both neuroimmune disease models and human patients, resulting in neuropathology. Further support for B cell function as APCs in OMAS comes from the increased frequency in OMAS of HLA-DOB*01, an HLA allele expressed predominantly in B cells that modulates presentation of immunodominant epitopes (reviewed in Welsh and Sadegh-Nasseri, 2020; Jiang et al 2019). Finally, the observation of B cell trafficking, TLS- promoting chemokines in OMAS support the central role of B cells in TLS prevalence and B cell- T cell interactions accompanying both positive tumor outcomes and neuropathology.

If indeed a single mechanism underlies both CNS pathology and anti-tumor immunity in OMAS neuroblastoma patients, then OMAS tumors (and indeed, tumors of other paraneoplastic disease associated with neuroimmunity) may offer a system in which to study the cellular basis of neuronal damage in the CNS, which cannot be addressed in living patients. It is still unclear whether the observed diversity and polyclonality of tumor infiltrate in OMAS arises because of lymphocytic influx from the periphery, which would be consistent with the dominance of public TCRs in tumor like their representation in peripheral blood. Specific predictions made in the current study, such as the properties of OMAS-associated TLS B cells and selected T cells in in tumor control, and the putative role of auto-reactive T cells in brain neuropathology in OMAS, should be addressed in future work, using freshly isolated and cell-sorted CSF and tumor samples and in humanized mouse models. Our work supports renewed focus on antigen-presenting B cells as potentiators of cancer immunotherapy, through generation of long-lasting tertiary lymphoid structures to promote tumor destruction. Modulation of accompanying autoimmunity will be a critical bottleneck for clinical applications.

## METHODS

### Patient tumor samples

We retrospectively procured all primary tumor samples (N=38) available from the COG ANBL00P3 clinical trial, in which the efficacy of IV immunoglobulins (Ivig) in neuroblastoma patients with OMAS was tested (de Alarcon et al 2018). All patients enrolled were <8 years old with biopsy-proven, newly diagnosed neuroblastoma and OMAS. Samples collected from each patient included tumor tissue, cerebrospinal fluid (CSF) and blood sera from time of diagnosis. We also sequenced 13 patients each with low-risk (LR) and high-risk (HR) non-OMAS neuroblastomas as comparators. We obtained reliable data from all samples, using the Illumina RNA Access platform, an exon capture kit designed to salvage usable data from low-quality RNA samples. However, as a consequence of using this platform, our ability to harmonize our data with existing neuroblastoma RNA-seq datasets (using other platforms) was rather limited.

### RNA sequencing

Patient tumor RNA was sequenced with 2 x 150 bp, paired end sequencing, using the TruSeq RNA Access kit from Illumina (now called TruSeq RNA Exome; Qiagen, Valencia CA, USA) and quantified on a NanoDrop spectrophotometer. RNA purity and integrity was assessed by Agilent 4200 Tapestation. RNA integrity (RIN) scores for the samples varied from 1 to 7.9, though all samples had DV200 values of >30%. Sequencing libraries were prepared from 100ng total RNA from each sample, and were run on high output flowcells on an Illumina NextSeq 500, yielding an average of 30M reads per sample. Paired-end sequence reads were analyzed according to currently available best practices for whole-transcriptome analysis, as described below.

### RNA-seq data analysis

Raw FASTQs from both the OMAS/LR/HR cohort and NCI TARGET (Pugh TJ et al 2013) datasets were processed using fastq-mcf (https://github.com/ExpressionAnalysis/ea-utils/blob/wiki/ FastqMcf.md):

http://expressionanalysis.github.io/ea-utils/; parameters: --max-ns 4 –qual-mean 25 -H -p 5 -q 7 - l 25). Clipping completely removed reads with large homopolymers, overall low base quality scores or less than 25 nucleotides and removes low quality bases at the end of the sequence and adapters. These clipped reads were aligned to the human reference genome hg19 using STAR v2.4 (Dobin et al 2013) and a UCSC reference transcriptome supplemented with lincRNAs from

Ensembl. RSEM v1.2.14 (https://github.com/ExpressionAnalysis/STAR-SEQR) was used for both gene and isoform quantification. RNA fusion events were detected using STAR-SEQR v0.6.5 (Ritchie et al 2014) parameters: -m 1).

Differential expression analysis was performed using Q2 Solutions’ ensemble two group comparisons suite. This method summarizes the differential expression p-values and classification probabilities from five tools—t-test, limma4, DESeq2 ((Ritchie et al 2014), edgeR (Anders and Huber, 2010) and EBSeq (Robinson MD, McCarthy DJ, Smyth GK 2010)—to produce a new p-value for differential expression. For any given gene, the p-values of each constituent model are input into a logistic regression model, which estimates the probability that the gene is differentially expressed. This probability is transformed into a p-value for differential expression by comparing it against its empirical cumulative distribution as estimated by bootstrap resampling of TCGA data from various cancer types.

### HLA Typing

HLA types were identified in both OMAS/LR/HR and TARGET datasets using the default parameters of HLAProfiler (Buchkovich ML et al. 2017) and each allele tested for enrichment. For some genes, HLAProfiler identified alleles in less than 25% of samples. Alleles from these genes or alleles identified in only a single sample were excluded from the enrichment analysis. Significance of enrichment was determined by testing the distribution of each allele among patient subgroups compared to all alleles for the gene in the population (n=2*number of samples). Fisher’s exact test p-values were adjusted for multiple hypothesis testing using a Benjamini-Hochberg correction. Significantly enriched alleles are shown in Table 1 and complete results are presented in Table S4.

### Immune landscape signatures

Immune landscape signatures, including cytotoxic lymphocyte activity (esp. CD8 T cells), B cell activity, IFNγ levels, T cell trafficking, immune suppression activity from myeloid-derived cells (M2TAM cells, TGFB1 levels, PD-L1, etc), checkpoint ratios, and stromal responses, were detected in each sample as described in (Jones WD et al 2020). These immune signature scores represent weighted averages of (log) expression levels of genes within each signature. Immune subtypes were tested for enrichment in OMAS patients using a Fisher’s exact test with correction for multiple testing using Benjamini-Hochberg. Features that show statistically significant differences between OMAS and non-OMAS samples were plotted in a separate box (top), and features not showing significant differences between groups were plotted in a heatmap below. Clustering of samples was performed according to maximize similarity of gene expression patterns in heatmap for significant features in upper box.

#### Variant Identifications

Raw FASTQ were processed with a pipeline optimized for variant calling in RNA-seq data. First, raw reads were processed using FASTP v0.19.4 (Chen S et al 2018) and the parameters: -- trim_poly_g, --trim_poly_x, --cut_by_quality3, --cut_mean_quality 20, --n_base_limit 4, -- qualified_quality_phred 15, --length_required 25, --complexity_threshold 30, -- low_complexity_filter, --correction, --html. Trimmed reads were aligned against the GRCh37 reference genome and GENCODE v27 transcriptome using the STAR v2.5.4b aligner (parameters: --runMode alignReads, --alignSJDBoverhangMin 2, --alignSJoverhangMin 8, -- chimFilter None, --chimJunctionOverhangMin 10, --chimMainSegmentMultNmax 10, -- chimOutType SeparateSAMold, --chimScoreDropMax 30, --chimScoreMin 1, -- chimScoreSeparation 7, --chimSegmentMin 10, --chimSegmentReadGapMax 3, -- outFilterIntronMotifs RemoveNoncanonicalUnannotated, --outFilterMultimapNmax 20, -- outFilterMultimapScoreRange 1, --outFilterScoreMinOverLread 0.66, --outMultimapperOrder Random, 0-outSAMstrandField intronMotif, --outSAMunmapped Within, --quantMode TranscriptomeSAM, --readFilesCommand zcat, --twopassMode, Basic). Variants were detected using “GATK best practices for variant calling on RNA-seq”, using Sentieon’s suite of tools (Freed F, Aldana R, Weber JA, Edwards JS BioRxiv) in place of GATK. Gene expression was evaluated in-pipeline using RSEM v1.3.0. These gene counts, as well as other metrics such as coverage statistics, gene region annotations, RNA editing sites, and clinVar and dbSNP annotations, were input as features into a random forest model which further filtered variants and removed false positive variant predictions. Genes containing exonic variants in one or more samples were examined for enrichment of these variants in OMAS patients. Significance was determined using Fisher’s exact test and corrected for multiple tests using Benjamini-Hochberg. Significant genes were further examined to identify any single variants driving the significance results. For each significant gene, SNPs within the gene were tested independently for enrichment in OMAS patients, with Benjamini-Hochberg correction for multiple tests.

### Immune Subtype Classifier

The Immune Subtype classifer, as described in (Thorsson et al 2018) and updated in Gibbs DL (BioRXiv) was applied to the RNA-seq data collected in the current study, as well as to previously published neuroblastoma data from TARGET (Pugh TJ et al 2013) and to data from the Pan Cancer Atlas (Hoadley KA et al 2018)

### TCR repertoire analysis

#### Tumor genomic DNA

Tumor genomic DNA was obtained from COG, and 31 OMAS, 13 LR and 13 HR patient samples were sequenced for TCRβ locus, using the Adaptive Biotechnologies Immunoseq platform. Since input genomic DNA samples were not of uniform concentration, to compute repertoire size, total number of sequence reads obtained were normalized for the amount of input DNA loaded into the sequencing assay.

#### Data cleaning and normalization

For repertoire analysis, CDR3β sequence reads that are in-frame and have no stop codon were considered; all other sequences were filtered out. For each amino acid sequence in a given sample, we summed the frequencies of all its nucleotide variants (due to convergent recombination) to obtain the frequency associated with the amino acid sequence in the given sample.

#### Data analysis

All computations were done using R (R version 3.6.3), running on a CentOS Linux 7 core. Data manipulation, plotting, and standard statistical tests were done by base R and standard packages. All computations involving, clonality, diversity and Horn similarity, were done using the same subsampling scheme. We subsampled all patient TCRβ repertoires to a common size (1,382 reads), computed the statistic and averaged the value of the statistic over 100 such iterations. Sampling was done by the sample function in base R. Shannon index and Horn similarity were computed using the vegan community ecology package (version 2.5-6). Average values over 100 subsampling iterations were plotted using ggplot, with the mean value for each patient group indicated with a red line. Unless otherwise indicated, comparisons between groups were made using Wilcoxon rank sum test, and FDR corrected for number of tests.

#### TCR Sharing Level

For each amino acid CDR3 sequence, we calculated its sharing level in the cohort, i.e., to how many samples it belongs. For each sharing level, we calculated the number of sequences that have this sharing level. Figure 4F describes in log-log scale the relative frequency of sequences in each sharing level. In Figure S3A, we compared the sharing level within neuroblastoma patient group to the sharing level in PBMC of healthy individuals as captured by the Emerson data set (Emerson et al 2017). Each sequence was plotted according to its Emerson sharing level (X axis) and Patient Group sharing level (OMAS/LR/HR; Y axis). Some of the sequences highlighted in color are given in the “Overshared” sequences in Table S6.

#### Emerson data set

To estimate background frequencies of TCRβ receptor sequences, we used the Emerson data set (Emerson RO et al 2017), a set of 786 patient repertoires healthy volunteers (666 bone marrow samples from one cohort, and 120 peripheral blood samples from a second cohort). The observed frequencies of the public TCRs in this study are concordant with computed probabilities based on recombination frequencies and selection from the lab of Alexandra Walczak. The Emerson Dataset data was downloaded from the Adaptive web site (https://www.adaptivebiotech.com/immuneACCESS DOI https://doi.org/10.21417/B7001Z ).

#### BCR repertoire analysis

For this analysis, we included available material from 37 OMAS-associated neuroblastomas, 13 LR, and 13 HR non-OMAS-associated neuroblastomas. IgH sequencing was performed on genomic DNA using the Adaptive Biotechnologies platform.

#### Data Analysis

For BCR analysis, we used the immcantation portal packages to compute gene usage, clonality, clustering, mutation frequency and diversity. All computations were done using R (R version 3.6.3), running on ubuntu 16. Data manipulation, plotting, and standard statistical tests were done using base R and standard packages. Diversity and Shannon index analysis was done using alakazam and shazam R packages from immcantation (Gupta NT and Vander Heiden JA et al 2015). Shannon index was subsampled to 219 sequences per sample. Clonality was performed using Change-O from immcantation. Unless otherwise indicated, comparisons between groups were made using Wilcoxon rank sum test, and FDR corrected for number of tests.

#### IgH Gene assignment

IgH sequences were aligned to IGHV, IGHD, and IGHJ genes by applying IgBlast (Ye J et al 2013) using a reference germline that was downloaded from IMGT in 2017. The repertoires were sequenced using the Adaptive Biotechnologies ImmunoSeq platform, which returns only a partial V 25ssignme. This can cause mis-assignment of the V gene. Thus, for better clone inference for each patient, clones were defined as the same V family, J gene, and junction length using Change-O (Gupta NT and Vander Heiden JA et al 2015 et al., 2015). The cutoff threshold was determined with the shazam package (Gupta NT and Vander Heiden JA et al 2015 et al., 2015).

#### IgH Clusters

To define clusters of sequences, all subjects’ repertoires were pooled, and clusters were inferred by the DefineClones function from Change-O using the complete linkage method. The clusters were defined as sequences that share the same V family, J gene, and junction length. We also required a minimum of 85% amino acid identity across the junction sequence for inclusion. Clusters containing at least one sequence from at least 7 OMAS subjects were chosen for plotting.

#### Diversity analysis

Diversity analysis, using Shannon diversity index, was performed using the alakazam package (Gupta et al., 2015), where each sample was subsampled 100 times to a minimum repertoire size (219 sequences) with sequence replacement. Significance was determined using the Wilcoxon test and p-values were corrected for multiple tests with FDR.

#### Mutation analysis

Mutation frequency of a sequence was calculated as the number of mutation compared to the V germline sequence devided by the length of the V region sequences. For each subject the sequences for each V family were grouped and the median mutation frequency was selected. Significance was determined using the Wilcoxon test and p-values were corrected for multiple tests with FDR.

#### IGHV gene usage

IgH sequences obtained using the ImmunoSeq platform carry only a partial V region, which hinders accurate 26ssignment of V gene identity. To avoid mis-assignment biases, uncertain or unreliable gene assignments were filtered out using the RAbHIT package (Peres at al., 2019). Then, relative gene usage was calculated using the alakazam package (Gupta et al., 2015). Significance was determined using the Wilcoxon test and p-values were corrected for multiple tests with FDR.

### XG Boost: building a binary classifier out of RNA-seq data

Machine learning procedures were carried out using the python scikit-learn (version 0.18.2) and XGBoost package. We chose Gradient Boosting Decision Trees (specifically eXtreme Gradient Boosting, XGBoost (Chen et al., 2016)) as the prediction algorithm for its ability to capture non-linear interactions between features, its efficiency and the fact that is has been successfully used in a wide range of applications.

Due to the relatively low number of samples available, we used leave-one-out as the cross-validation scheme and did not perform hyperparameter optimization to avoid reducing the sample size even further by putting aside a dedicated subset used only for model optimization. For each iteration, XGBClassifier was trained on FPKM values from all but one sample, and the resulting

model was used to predict the class of the left out sample (either OMAS vs non-OMAS, OMAS vs HR, or OMAS vs LR). The performance was scored using the area under the ROC curve as a metric. ROC curves for each comparison, as well as top features for each XGBoost model, are given in Figure S1. Feature importance and effect on the model was determined using SHAP analysis (Lundberg et al., 2020).

#### Immunohistochemistry, TLS imaging, and histological scoring

Paraffin embedded sections from OMAS and non-OMAS patient tumors (5 micron sections, charged slides, air dried) were obtained from primary tumor resection (with two exceptions, which were optained from biopsies). Sections were obtained on slides from Children’s Oncology Group. Images of H&E stained sections from the same specimens, which had been prepared, stained using standard methods, and imaged previously by COG at 40X magnification, were also obtained for scoring.

#### Immunohistochemistry

Unstained slides of formalin-fixed, paraffin-embedded sections were stained as follows: Slides were rinsed in 2 changes of xylene for 5 min each, then rehydrated in a series of descending concentrations of ethanol. Slides were then treated in a pressure cooker with antigen unmasking solution (Vector Laboratories H-3300) for 30 minutes. After cooling, slides were rinsed in 0.1M Tris Buffer, and then blocked in 0.1M Tris buffer, 0.01% tween with 2% fetal bovine serum for 15 min. For primary antigen detection, the following primary antibody combinations were used: a) Rabbit anti-CD3 (1:50, Dako A0452), incubated overnight, and mouse anti-CD20 (1:500, Dako M0755), which were incubated with the slides for 1 hour at room temperature, and b) Goat anti-human CD4 (1:400, R&D Systems AF-379-NA) and Rabbit anti-human CD8 (1:400, Thermo RB-9009-P0), which were both incubated for 1hr at room temperature. After primary antibody staining, slides were again rinsed several times in 0.1M Tris Buffer with 0.01% Tween, and then incubated with the following secondary antibody combinations: For CD3/CD20 detection, Alexa 488 anti-Rabbit (Life Technologies, A21206), with Alexa 594 Anti-Mouse (Life Technologies, A11032) were used. For CD4/ CD8 detection, Alexa 488 anti-Goat (Life Technologies, A11055) with Alexa 594 anti-Rabbit (Life Technologies, A21207) at a 1:400 dilution were used. All slides were incubated with secondary antibodies for 1hr at room temp. Slides were rinsed several times in 0.1M Tris/0.01% Tween, then counterstained for 5min in DAPI Hydrochloride (Sigma 32670). Slides were then rinsed, and then coverslipped with Prolong Gold (Life Technologies, P36930). Slides were digitally scanned at 20x magnification (Aperio IF, Leica Biosystems).

**For Ki67 staining**, coverslips from stained slides were removed by incubating the slides in 1xPBS at 37°C overnight, and then washed for 2 hours in 1x PBST with several changes, before proceeding to Ki67 staining. Without removing prior staining (CD3-alexa 488/CD20 alexa 594/DAPI), slides were further stained using Rabbit monoclonal anti-Ki-67:Alexafluor 647 direct conjugate (Abcam ab196907, 1:100) at 4°C overnight. Slides were then washed in 1x PBST with several changes for 2 hours before mounting and coverslipping in Slow-fade Gold mounting medium (ThermoFisher).

### Histological and immunohistochemical examination of tumor specimens

Formalin-fixed paraffin-embedded tissue sections stained with hematoxylin and eosin (HE) from each of the tumor samples included in the study were histologically revised to confirm the initial diagnosis of neuroblastomas or ganglioneuroblastomas applying the criteria for classification of neuroblastic tumors suggested by the International Neuroblastoma Pathology Committee (Shimada H et al 1999). Signs of differentiation tendency in the neuroblastic tumors, such as presence of neuropils, Homer-Wright rosettes, and different stages of maturation towards ganglion cells were recorded. Additionally, we assessed the possible presence of tertiary lymphoid structures containing lymphatic follicles with or without germinal centers according to previously published quantification criteria (none= 0; present in <10% of tumor tissue = 1+; present in 10% to 50% of tumor tissue = 2+; present in >50% of tumor tissue = 3+; Hudlebusch et al, 2011).

Lymphocytic populations in the tumor-associated lymphoid structures and elsewhere in the tumors were assessed by immunofluorescent staining of tissue sections using primary antibodies against human CD20 and CD3, a B-cell and T-cell marker, respectively, as described above. Proliferation activity in the germinal centers of lymphatic follicles was assessed using immunofluorescent staining against human Ki67, as described above.

#### Imaging of TIL immunohistochemistry

Images were acquired on a Leica LMD upright widefield microscope driven by the LAS X acquisition software, with a 20X objective. Raw images were identically scaled and then exported as TIFFs.

## Acknowledgments

This manuscript is dedicated in memory of Jessica A. Panzer, a remarkable physician-scientist and Nir Friedman, a rare and brilliant computational immunologist, both wonderful human beings who contributed to this work but did not live to see its completion. This work was inspired by OMAS/neuroblastoma patient and survivor, MDB, and by neuroblastoma patient, Yazan El Kooka, who is with us still in spirit.

This work was funded by grants to MIR and JAP from the Pablove Foundation, the Rally Foundation, Open Hands/Overflowing Hearts Foundation, and The Truth 365 Foundation; by NIH grant R35 CA220500 (J.M.M.), the Giulio D’Angio Endowed Chair (J.M.M.) and RC1MD004418 to the NCI-TARGET consortium (COG SDC grant U10 CA180899), and NCTN Operations Center Grant U10CA180886 and NCTN Statistics & Data Center Grant U10CA180899, and support from St. Baldrick’s Foundation to Children’s Oncology Group. MIR would also like to thank Naveen and Crystal Viswanatha, for creating and funding the dedicated Pablove OMAS research grant, Anne Spurkland and Vlad Vigdorovich for thoughtful, critical reading of the manuscript, Paolo Fortina, David Lynch, Daniel Martinez, Phil Bradley, Avi Jacob, and Matthew Weitzman for generous technical advice and effort, and Stephen J. Tapscott for sage guidance and unwavering support. Finally, we thank all patient families.

## Disclaimer

The content is solely the responsibility of the authors and does not necessarily represent the official views of the National Institutes of Health

## Author Contributions

MIR designed the project with JAP and led the project, performed tumor imaging, data analysis and synthesis, wrote the manuscript with contributions from all authors; EG led TCR analysis with contributions from DR and supervision by NF; MB carried out bioinformatics analysis including differential expression, SNP analysis, immune landscape signature and HLA analyses, and implementation of immune subtype pipeline; MM carried out machine learning experiments (XGBoost), and contributed statistical and bioinformatic analysis; AP led BCR analysis under the supervision of GY; ES-R carried out the histopathological scoring and immunohistochemical phenoypting; AS contributed to immunohistochemical analyses; DLG carried out additional immune subtype analysis and bioinformatics; IU contributed to bioinformatic analysis; AN, MI and PdA led the clinical trial from which samples for this study were obtained; VW supervised transcriptome analysis, HLA analysis and design, provided bioinformatics support; JMM supervised the project.

## Supplementary Figure Legends

**Figure S1. A machine learning classifier, XGBoost, identifies gene expression signatures that distinguish OMAS neuroblastoma from non-OMAS neuroblastoma. (A-C) Shap plots for signatures of models distinguishing OMAS neuroblastoma from control neuroblastoma.** Normalized, transcriptome-wide expression was compared for all samples in comparison groups, except one sample set aside for model validation (using Leave-one-out). For each model, SHAP value (a score indicating feature importance for model; Lundberg et al 2020) is indicated on the X axis, gene features are given on the Y axis. Individual patients represented as dots, gene expression value for each feature given as a color (range at right). Pink=high expression, blue=low expression. Distance from x=0 indicates contribution of gene feature to model. **(A)** Top twenty gene features of model distinguishing OMAS neuroblastoma from non-OMAS neuroblastoma. **(B)** Top twenty gene features of model distinguishing OMAS from HR non-OMAS neuroblastoma. **(C)** Top twenty features of model distinguishing OMAS neuroblastoma from LR non-OMAS neuroblastoma. **(D-F) auROC curves for each XGB model.** For each model in (A-C), performance was scored using the area under the ROC curve as a metric. auROC=1 indicates 100% prediction accuracy for classification of left out sample. For each curve, true postive rate (Y-axis) is plotted against corresponding false positive rate (X-axis). Blue curve= XGB model, grey dotted line= neutral model. **(D)** auROC curve for OMAS vs non-OMAS model. **(E)** auROC curve for OMAS vs HR non-OMAS model. **(F)** auROC curve for OMAS vs LR non- OMAS model. **(G)** Top ten single gene features driving OMAS vs non-OMAS neuroblastoma model from XGBoost. For each gene, a box plot is given of FPKM values (Y axis) for the gene features in each patient (scatter), with median value indicated for each patient group. OMAS= red, non-OMAS=green.

**Figure S2. Correlation of single gene expression with neurological symptom severity scores nominates candidate OMAS autoantigens.** Correlation of expression values (normalized RPKM) for single gene features with neurological severity score (range 0-14) was tested using Spearman correlation. **(A)** Table of genes with significant correlation of expression with neurological severity score. For each gene, gene, Spearman correlation (R), p value, gene name, gene ID, and gene function (via Genecards) are given. Genes whose expression in OMAS is negatively correlated with severity score are highlighted in red. For genes with positive correlation of gene expression with symptom severity: Green boxes= neuronal cell surface receptors/channels, dark blue boxes= cell adhesion molecules. **(B-D)** Single candidate gene plots of gene expression level (RPKM) as a function of symptom severity score. **(B)** NCAN. **(C)** HTR6. **(D)** ADRA2C.

**Figure S3. TCR sharing and clonal structures of patient TCR repertoires in this study. (A)** TCR sharing levels of TIL TCRβ sequences compared to sharing in Emerson data set. For each patient group, sharing level of individual TCRs (dots) are plotted according to their within-group sharing level (Y axis) and their Emerson sharing level (X axis; range 0-786). Colored dots indicate sequences that are more highly shared within their group than within the Emerson dataset (“overshared”). **(B)** Clonal structures of TCR repertoires for each patient. Clonal frequencies for the top 1 (dark orange), top 10 (green), top 50 (light blue) and top 100 (light orange) TCRs in each patient repertoire were summed, and plotted as a stacked bar to fraction of the repertoire occupied by each clonal subset. Samples are plotted along X axis, with stacked bars for summed frequencies within repertoire plotted on Y axis. **(C)** Sharing levels of TCRs in each patient repertoire compared to Emerson sharing levels. For each patient, repertoires are represented as stacked bars indicating the fraction of each patient repertoire that is shared by patients in the Emerson dataset. Dark blue indicates sequences not represented in Emerson (private sequences). Yellow= shared by <25% of patients in Emerson; Green= shared by 25-50% of patients in Emerson; Light blue= shared by 50-75% of patients in Emerson; Orange=share by 75-100% of patients in Emerson.

**Figure S4. BCR Repertoires of OMAS-associated neuroblastoma are largely similar to non-OMAS neuroblastoma in IGH gene usage and other junction features.** For all plots shown, samples are color coded according to patient group: Green=OMAS, orange= LR, purple= HR. Combinations whose differential usage between groups have FDRq<0.05 indicated with a *; orange star indicates HR-LR is significant, dark blue star indicates OMAS-LR is significant, brown star indicates OMAS-HR combination is significant. **(A)** IGHV gene usage. Box plot showing observed frequencies for each gene (X axis) for each patient (dots) and within each group. **(B)** IGHJ gene usage. Box plot showing observed frequencies for each gene (X axis) for each patient (dots) and within each group. **(C)** V gene family-J gene family usage. V gene-J gene family combinations were scored for observed frequencies in each patient group. Top 30 combinations are plotted. **(D)** Junction length. Violin plot of observed junction lengths for all BCRs in each patient repertoire. **(E)** V gene family-J gene family-Junction length. VJ-Junction length combinations were scored for observed frequencies in each patient group.

**Figure S5. Additional immunohistochemical study of tertiary lymphoid structures in OMAS neuroblastomas.** For panels (A-C), B cells are labeled with anti-CD20 antibody (red) and CD8+ T cells are labeled in green. **(A)** Tertiary lymphoid structures in OMAS tumors. **(B)** Nuclear Ki67 staining of proliferating cells in and around tertiary lymphoid structures in OMAS tumors. Arrows indicate nuclei labeled with anti-Ki67 antibody (purple); membrane associated staining was scored as background. **(C)** Localization of tumor infiltrating lymphocytes to spindle-like processes resembling neuronal processes. **(D)** Summary and quantification of tumor histopathology findings. Presence of tertiary lymphoid structures and germinal centers were scored utilizing a previously published scale (Hudlebusch, et al 2011), as follows: in none = 0; present in <10% of tumor tissue = 1+; present in 10% to 50% of tumor tissue = 2+; present in >50% of tumor tissue = 3+). Patient group is indicated by color: red= HR non-OMAS; yellow= LR non-OMAS; green= OMAS.

## Supplementary Table Legends

**Table S1. Differential gene expression analysis.** Sheet one (samplesOMSnonOMS) lists the samples in the comparison. Sheet two (OMSnonOMS_genes) lists features of differential expression analysis, filtered for FDRq<0.05. Samples colored in red have Log2(FC)≥1, and were used as input for ENRICHR (Figure 1 panel B). Samples colored in blue have Log2(FC)≥-1, and were used as input for ENRICHR (Figure 1 panel C). Sheet three (OMSnonOMS_genes.support) provides additional information to support values given in sheet 2. Sheet four (samplesLROMS) lists the samples in the comparison. Sheet five (LROMS_genes) lists features of differential expression analysis, filtered for FDRq<0.05. Samples colored in red have Log2(FC)≥1; samples colored in blue have Log2(FC)≥-1. Sheet six (LROMS_ genes.support) provides additional information to support values given in sheet 5.

**Table S2. Complete SNP burden enrichment by gene, and complete table of HLA allele enrichment in OMAS in this study comparing OMAS tumors in this study to LR and HR non-OMAS controls from this study and TARGET.** P values and adjusted P values <0.05 are labeled in yellow.

**Table S3: TCR sharing.** Sheets 1-3: TCRdist results. Output of TCRdist (Dash P et al 2017) for all TCR repertoires sequenced in the current study (OMAS=31, LR=13, HR=13). For TCRdist100, results are given for similarity search using only the top 100 clones in each patient repertoire. TCRdist1000 indicates results using top 1000 clones in each patient repertoire. TCRdistALL contains output using all TCRs in each patient repertoire for comparison. P values were adjusted using FDR correction for number of tests. Sheet 4: Over-shared TCRs. For each TCR sequence, sharing level is defined as number of individuals in the patient group with that sequence in their repertoire. For TCR repertoires, the total number of patients in each cohort are: OMAS=31, LR=13, HR=13.

## REFERENCES

1. Altman, A.J., and Baehner, R.L. (1976). Favorable prognosis for survival in children with coincident opso-myoclonus and neuroblastoma. Cancer 37, 846–852.

2. Bagaeva, L.V., Rao, P., Powers, J.M., and Segal, B.M. (2006). CXC chemokine ligand 13 plays a role in experimental autoimmune encephalomyelitis. J Immunol 176, 7676–7685. 10.4049/jimmunol.176.12.7676.

3. Bernards, R., Dessain, S.K., and Weinberg, R.A. (1986). N-myc amplification causes down-modulation of MHC class I antigen expression in neuroblastoma. Cell 47, 667–674. 10.1016/0092-8674(86)90509-x.

4. Berridge, G., Menassa, D.A., Moloney, T., Waters, P.J., Welding, I., Thomsen, S., Zuberi, S., Fischer, R., Aricescu, A.R., Pike, M., et al. (2018). Glutamate receptor delta2 serum antibodies in pediatric opsoclonus myoclonus ataxia syndrome. Neurology 91, e714–e723. 10.1212/WNL.0000000000006035.

5. Brady, S.W., Liu, Y., Ma, X., Gout, A.M., Hagiwara, K., Zhou, X., Wang, J., Macias, M., Chen, X., Easton, J., et al. (2020). Pan-neuroblastoma analysis reveals age- and signature-associated driver alterations. Nature communications 11, 5183. 10.1038/s41467-020-18987-4.

6. Buchkovich, M.L., Brown, C.C., Robasky, K., Chai, S., Westfall, S., Vincent, B.G., Weimer, E.T., and Powers, J.G. (2017). HLAProfiler utilizes k-mer profiles to improve HLA calling accuracy for rare and common alleles in RNA-seq data. Genome medicine 9, 86. 10.1186/s13073-017-0473-6.

7. Bugatti, S., Manzo, A., Vitolo, B., Benaglio, F., Binda, E., Scarabelli, M., Humby, F., Caporali, R., Pitzalis, C., and Montecucco, C. (2014). High expression levels of the B cell chemoattractant CXCL13 in rheumatoid synovium are a marker of severe disease. Rheumatology (Oxford) 53, 1886–1895. 10.1093/rheumatology/keu163.

8. Byrne, K.T., and Turk, M.J. (2011). New perspectives on the role of vitiligo in immune responses to melanoma. Oncotarget 2, 684–694. 10.18632/oncotarget.323.

9. Cabrita, R., Lauss, M., Sanna, A., Donia, M., Skaarup Larsen, M., Mitra, S., Johansson, I., Phung, B., Harbst, K., Vallon-Christersson, J., et al. (2020). Tertiary lymphoid structures improve immunotherapy and survival in melanoma. Nature 577, 561–565. 10.1038/s41586-019-1914-8.

10. Carlson, C.S., Emerson, R.O., Sherwood, A.M., Desmarais, C., Chung, M.W., Parsons, J.M., Steen, M.S., LaMadrid-Herrmannsfeldt, M.A., Williamson, D.W., Livingston, R.J., et al. (2013). Using synthetic templates to design an unbiased multiplex PCR assay. Nature communications 4, 2680. 10.1038/ncomms3680.

11. Chen, E.Y., Tan, C.M., Kou, Y., Duan, Q., Wang, Z., Meirelles, G.V., Clark, N.R., and Ma’ayan, A. (2013). Enrichr: interactive and collaborative HTML5 gene list enrichment analysis tool. BMC bioinformatics 14, 128. 10.1186/1471-2105-14-128.

12. Chen, F., Zhang, Y., Varambally, S., and Creighton, C.J. (2019). Molecular Correlates of Metastasis by Systematic Pan-Cancer Analysis Across The Cancer Genome Atlas. Molecular cancer research : MCR 17, 476–487. 10.1158/1541-7786.MCR-18-0601.

13. Chen, T., Guestrin, C. (2016). XGBoost: reliable large-scale tree boosting system. .held in San Francisco, CA, pp. 13–17.

14. Cooper, R., Khakoo, Y., Matthay, K.K., Lukens, J.N., Seeger, R.C., Stram, D.O., Gerbing, R.B., Nakagawa, A., and Shimada, H. (2001). Opsoclonus-myoclonus-ataxia syndrome in neuroblastoma: histopathologic features-a report from the Children’s Cancer Group. Medical and pediatric oncology 36, 623–629. 10.1002/mpo.1139.

15. Dalmau, J., Tüzün, E., Wu, H-y., Masjuan, J., Rossi, J.E., Voloschin, A., Baehring, J.M., Shimazaki, H., Koide, R., King, D., Mason, W., Sansing, L.H., Dicheter, M.A., Rosenfeld, M.R., and Lynch, D.R. (2007) Paraneoplastic anti-N-methyl-D-aspartate receptor encephalitis associated with ovarian teratoma. Annals of Neurology 61, 25–36. 10.1002/ana.21050.

16. Dalmau, J., Armangue, T., Planaguma, J., Radosevic, M., Mannara, F., Leypoldt, F., Geis, C., Lancaster, E., Titulaer, M.J., Rosenfeld, M.R., and Graus, F. (2019). An update on anti-NMDA receptor encephalitis for neurologists and psychiatrists: mechanisms and models. Lancet neurology 18, 1045–1057. 10.1016/S1474-4422(19)30244-3.

17. Darnell, R.B., and Posner, J.B. (2003). Paraneoplastic syndromes involving the nervous system. The New England journal of medicine 349, 1543–1554. 10.1056/NEJMra023009.

18. Dash, P., Fiore-Gartland, A.J., Hertz, T., Wang, G.C., Sharma, S., Souquette, A., Crawford, J.C., Clemens, E.B., Nguyen, T.H.O., Kedzierska, K., et al. (2017). Quantifiable predictive features define epitope-specific T cell receptor repertoires. Nature 547, 89–93. 10.1038/nature22383.

19. De Grandis, E., Parodi, S., Conte, M., Angelini, P., Battaglia, F., Gandolfo, C., Pessagno, A., Pistoia, V., Mitchell, W.G., Pike, M., et al. (2009). Long-term follow-up of neuroblastoma-associated opsoclonus-myoclonus-ataxia syndrome. Neuropediatrics 40, 103–111. 10.1055/s-0029-1237723.

20. De Mattos-Arruda, L., Sammut, S.J., Ross, E.M., Bashford-Rogers, R., Greenstein, E., Markus, H., Morganella, S., Teng, Y., Maruvka, Y., Pereira, B., et al. (2019). The Genomic and Immune Landscapes of Lethal Metastatic Breast Cancer. Cell reports 27, 2690–2708 e2610. 10.1016/j.celrep.2019.04.098.

21. Dendrou, C.A., Petersen, J., Rossjohn, J., and Fugger, L. (2018). HLA variation and disease. Nat Rev Immunol 18, 325–339. 10.1038/nri.2017.143.

22. Dieu-Nosjean, M.C., Giraldo, N.A., Kaplon, H., Germain, C., Fridman, W.H., and Sautes-Fridman, C. (2016). Tertiary lymphoid structures, drivers of the anti-tumor responses in human cancers. Immunological reviews 271, 260–275. 10.1111/imr.12405.

23. Emerson, R.O., DeWitt, W.S., Vignali, M., Gravley, J., Hu, J.K., Osborne, E.J., Desmarais, C., Klinger, M., Carlson, C.S., Hansen, J.A., et al. (2017). Immunosequencing identifies signatures of cytomegalovirus exposure history and HLA-mediated effects on the T cell repertoire. Nature genetics 49, 659–665. 10.1038/ng.3822.

24. Ehrlich P. Ueber den jetzigen Stand der Karzinomforschung. Ned Tijdschr Geneeskd. 1909;5:273–290

25. Escobar, G., Mangani, D., and Anderson, A.C. (2020). T cell factor 1: A master regulator of the T cell response in disease. Sci Immunol 5. 10.1126/sciimmunol.abb9726.

26. Fukushima, H., Inoue, T., Takama, Y., Ishii, N., Okuno, T., Kobayashi, Y., Yoneda, A., Nakamura, T., Kuki, I., and Hara, J. (2017). Clinicopathological features of neuroblastic tumors with opsoclonus-myoclonus-ataxia syndrome: Follicular structure predicts a better neurological outcome. Pathology international 67, 503–509. 10.1111/pin.12591.

27. Gambini, C., Conte, M., Bernini, G., Angelini, P., Pession, A., Paolucci, P., Donfrancesco, A., Veneselli, E., Mazzocco, K., Tonini, G.P., et al. (2003). Neuroblastic tumors associated with opsoclonus-myoclonus syndrome: histological, immunohistochemical and molecular features of 15 Italian cases. Virchows Archiv : an international journal of pathology 442, 555–562. 10.1007/s00428-002-0747-1.

28. Giles, J.R., Kashgarian, M., Koni, P.A., and Shlomchik, M.J. (2015). B Cell-Specific MHC Class II Deletion Reveals Multiple Nonredundant Roles for B Cell Antigen Presentation in Murine Lupus. J Immunol 195, 2571–2579. 10.4049/jimmunol.1500792.

29. Graus, F., Delattre, J.Y., Antoine, J.C., Dalmau, J., Giometto, B., Grisold, W., Honnorat, J., Smitt, P.S., Vedeler, C., Verschuuren, J.J., et al. (2004). Recommended diagnostic criteria for paraneoplastic neurological syndromes. Journal of neurology, neurosurgery, and psychiatry 75, 1135–1140. 10.1136/jnnp.2003.034447.

30. Helmink, B.A., Reddy, S.M., Gao, J., Zhang, S., Basar, R., Thakur, R., Yizhak, K., Sade-Feldman, M., Blando, J., Han, G., et al. (2020). B cells and tertiary lymphoid structures promote immunotherapy response. Nature 577, 549–555. 10.1038/s41586-019-1922-8.

31. Hero, B., Clement, N., Ora, I., Pierron, G., Lapouble, E., Theissen, J., Pasqualini, C., Valteau-Couanet, D., Plantaz, D., Michon, J., et al. (2018). Genomic Profiles of Neuroblastoma Associated With Opsoclonus Myoclonus Syndrome. Journal of pediatric hematology/oncology 40, 93–98. 10.1097/MPH.0000000000000976.

32. Hero, B., Radojska, S., and Gathof, B. (2005). Opsomyoclonus Syndrome in infancy with or without neuroblastoma is associated with HLA-DRB1*01. In 4. held in Vancouver, Canada, (Pediatr Blood Cancer), pp. Pages 365–608.

33. Hudlebusch, H.R., Skotte, J., Santoni-Rugiu, E., Zimling, Z.G., Lees, M.J., Simon, R., Sauter, G., Rota,R., De Ioris, M.A., Quarto, M., Johansen, J.V., Jørgensen, M., Rechnitzer, C., Maroun, L.L., Schrøder, H., Petersen, B.L., and Helin K. (2011). MMSET Is Highly Expressed and Associated with Aggressiveness in Neuroblastoma. Cancer Research 71: 4226–35. 10.1158/0008-5472.CAN-10-3810

34. Ilicic, T., Kim, J.K., Kolodziejczyk, A.A., Bagger, F.O., McCarthy, D.J., Marioni, J.C., and Teichmann, S.A. (2016). Classification of low quality cells from single-cell RNA-seq data. Genome Biol 17, 29. 10.1186/s13059-016-0888-1.

35. Jiang, W., Adler, L.N., Macmillan, H., and Mellins, E.D. (2019). Synergy between B cell receptor/antigen uptake and MHCII peptide editing relies on HLA-DO tuning. Scientific reports 9, 13877. 10.1038/s41598-019-50455-y.

36. Jones, B.E., Tovar, K.R., Goehring, A., Jalali-Yazdi, F., Okada, N.J., Gouaux, E., and Westbrook, G.L. (2019) Autoimmune receptor encephalitis in mice induced by active immunization with conformationally stabilized holoreceptors. Science Translational Medicine 11:eaaw0044. 10.1126/scitranslmed.aaw0044

37. Jones, G.W., and Jones, S.A. (2016). Ectopic lymphoid follicles: inducible centres for generating antigen-specific immune responses within tissues. Immunology 147, 141–151. 10.1111/imm.12554.

38. Karlsson, L., Surh, C.D., Sprent, J., and Peterson, P.A. (1991). A novel class II MHC molecule with unusual tissue distribution. Nature 351, 485–488. 10.1038/351485a0.

39. Kazanietz, M.G., Durando, M., and Cooke, M. (2019). CXCL13 and Its Receptor CXCR5 in Cancer: Inflammation, Immune Response, and Beyond. Front Endocrinol (Lausanne) 10, 471. 10.3389/fendo.2019.00471.

40. Kinsbourne, M. (1962). Myoclonic encephalopathy of infants. Journal of neurology, neurosurgery, and psychiatry 25, 271–276.

41. Kulik, L., Laskowski, J., Renner, B., Woolaver, R., Zhang, L., Lyubchenko, T., You, Z., Thurman, J.M., and Holers, V.M. (2019). Targeting the Immune Complex-Bound Complement C3d Ligand as a Novel Therapy for Lupus. J Immunol 203, 3136–3147. 10.4049/jimmunol.1900620.

42. Magliozzi, R., Howell, O., Vora, A., Serafini, B., Nicholas, R., Puopolo, M., Reynolds, R., and Aloisi, F. (2007). Meningeal B-cell follicles in secondary progressive multiple sclerosis associate with early onset of disease and severe cortical pathology. Brain : a journal of neurology 130, 1089–1104. 10.1093/brain/awm038.

43. McGovern, V.J. (1975). Spontaneous regression of melanoma. Pathology 7, 91–99. 10.3109/00313027509092702.

44. Molnarfi, N., Schulze-Topphoff, U., Weber, M.S., Patarroyo, J.C., Prod’homme, T., Varrin-Doyer, M., Shetty, A., Linington, C., Slavin, A.J., Hidalgo, J., et al. (2013). MHC class II-dependent B cell APC function is required for induction of CNS autoimmunity independent of myelin-specific antibodies. The Journal of experimental medicine 210, 2921–2937. 10.1084/jem.20130699.

45. Moschovakis, G.L., Bubke, A., Friedrichsen, M., Falk, C.S., Feederle, R., and Forster, R. (2017). T cell specific Cxcr5 deficiency prevents rheumatoid arthritis. Scientific reports 7, 8933. 10.1038/s41598-017-08935-6.

46. Nelson, B.H. (2010). CD20+ B cells: the other tumor-infiltrating lymphocytes. J Immunol 185, 4977–4982. 10.4049/jimmunol.1001323.

47. Newman, A.M., Liu, C.L., Green, M.R., Gentles, A.J., Feng, W., Xu, Y., Hoang, C.D., Diehn, M., and Alizadeh, A.A. (2015). Robust enumeration of cell subsets from tissue expression profiles. Nature methods 12, 453–457. 10.1038/nmeth.3337.

48. Nordlund, J.J., Kirkwood, J.M., Forget, B.M., Milton, G., Albert, D.M., and Lerner, A.B. (1983). Vitiligo in patients with metastatic melanoma: a good prognostic sign. Journal of the American Academy of Dermatology 9, 689–696. 10.1016/s0190-9622(83)70182-9.

49. Peng, Y., Xiao, L., Rong, H., Ou, Z., Cai, T., Liu, N., Li, B., Zhang, L., Wu, F., Lan, T., et al. (2021). Single-cell profiling of tumor-infiltrating TCF1/TCF7(+) T cells reveals a T lymphocyte subset associated with tertiary lymphoid structures/organs and a superior prognosis in oral cancer. Oral Oncol 119, 105348. 10.1016/j.oraloncology.2021.105348.

50. Pipi, E., Nayar, S., Gardner, D.H., Colafrancesco, S., Smith, C., and Barone, F. (2018). Tertiary Lymphoid Structures: Autoimmunity Goes Local. Frontiers in immunology 9, 1952. 10.3389/fimmu.2018.01952.

51. Pranzatelli, M.R., Tate, E.D., McGee, N.R., Travelstead, A.L., Ransohoff, R.M., Ness, J.M., and Colliver, J.A. (2012). Key role of CXCL13/CXCR5 axis for cerebrospinal fluid B cell recruitment in pediatric OMS. Journal of neuroimmunology 243, 81–88. 10.1016/j.jneuroim.2011.12.014.

52. Pranzatelli, M.R., Tate, E.D., Travelstead, A.L., Barbosa, J., Bergamini, R.A., Civitello, L., Franz, D.N., Greffe, B.S., Hanson, R.D., Hurwitz, C.A., et al. (2006). Rituximab (anti- CD20) adjunctive therapy for opsoclonus-myoclonus syndrome. Journal of pediatric hematology/oncology 28, 585–593. 10.1097/01.mph.0000212991.64435.f0.

53. Pugh, T.J., Morozova, O., Attiyeh, E.F., Asgharzadeh, S., Wei, J.S., Auclair, D., Carter, S.L., Cibulskis, K., Hanna, M., Kiezun, A., et al. (2013). The genetic landscape of high-risk neuroblastoma. Nature genetics 45, 279–284. 10.1038/ng.2529.

54. Robins, H.S., Campregher, P.V., Srivastava, S.K., Wacher, A., Turtle, C.J., Kahsai, O., Riddell, S.R., Warren, E.H., and Carlson, C.S. (2009). Comprehensive assessment of T-cell receptor beta-chain diversity in alphabeta T cells. Blood 114, 4099–4107. 10.1182/blood-2009-04-217604.

55. Ruffin, A.T., Cillo, A.R., Tabib, T., Liu, A., Onkar, S., Kunning, S.R., Lampenfeld, C., Atiya, H.I., Abecassis, I., Kurten, C.H.L., et al. (2021). B cell signatures and tertiary lymphoid structures contribute to outcome in head and neck squamous cell carcinoma. Nature communications 12, 3349. 10.1038/s41467-021-23355-x.

56. Sansing, L.H., Tuzun, E., Ko, M.W., Baccon, J., Lynch, D.R., and Dalmau, J. (2007). A patient with encephalitis associated with NMDA receptor antibodies. Nature clinical practice. Neurology 3, 291–296. 10.1038/ncpneuro0493.

57. Sautes-Fridman, C., Petitprez, F., Calderaro, J., and Fridman, W.H. (2019). Tertiary lymphoid structures in the era of cancer immunotherapy. Nature reviews. Cancer 19, 307–325. 10.1038/s41568-019-0144-6.

58. Schuierer, S., Carbone, W., Knehr, J., Petitjean, V., Fernandez, A., Sultan, M., and Roma, G. (2017). A comprehensive assessment of RNA-seq protocols for degraded and low-quantity samples. BMC Genomics 18, 442. 10.1186/s12864-017-3827-y.

59. Shen, K., Xu, Y., Guan, H., Zhong, W., Chen, M., Zhao, J., Li, L., and Wang, M. (2018). Paraneoplastic limbic encephalitis associated with lung cancer. Scientific reports 8, 6792. 10.1038/s41598-018-25294-y.

60. Shlomchik, M.J., and Weisel, F. (2012). Germinal center selection and the development of memory B and plasma cells. Immunological reviews 247, 52–63. 10.1111/j.1600-065X.2012.01124.x.

61. Smith, J.L., Jr., and Stehlin, J.S., Jr. (1965). Spontaneous regression of primary malignant melanomas with regional metastases. Cancer 18, 1399–1415. 10.1002/1097-0142(196511)18:11<1399::aid-cncr2820181104>3.0.co;2-r.

62. Su, Z., Kishida, S., Tsubota, S., Sakamoto, K., Cao, D., Kiyonari, S., Ohira, M., Kamijo, T., Narita, A., Xu, Y., et al. (2017). Neurocan, an extracellular chondroitin sulfate proteoglycan, stimulates neuroblastoma cells to promote malignant phenotypes. Oncotarget 8, 106296–106310. 10.18632/oncotarget.22435.

63. Swann, J.B., and Smyth, M.J. (2007). Immune surveillance of tumors. The Journal of clinical investigation 117, 1137–1146. 10.1172/JCI31405.

64. Thorsson, V., Gibbs, D.L., Brown, S.D., Wolf, D., Bortone, D.S., Ou Yang, T.H., Porta-Pardo, E., Gao, G.F., Plaisier, C.L., Eddy, J.A., et al. (2018). The Immune Landscape of Cancer. Immunity 48, 812–830 e814. 10.1016/j.immuni.2018.03.023.

65. Trub, M., and Zippelius, A. (2021). Tertiary Lymphoid Structures as a Predictive Biomarker of Response to Cancer Immunotherapies. Frontiers in immunology 12, 674565. 10.3389/fimmu.2021.674565.

66. Valencia-Sanchez, C., and Zekeridou, A. (2021). Paraneoplastic Neurological Syndromes and Beyond Emerging With the Introduction of Immune Checkpoint Inhibitor Cancer Immunotherapy. Front Neurol 12, 642800. 10.3389/fneur.2021.642800.

67. Valpione, S., Mundra, P.A., Galvani, E., Campana, L.G., Lorigan, P., De Rosa, F., Gupta, A., Weightman, J., Mills, S., Dhomen, N., and Marais, R. (2021). The T cell receptor repertoire of tumor infiltrating T cells is predictive and prognostic for cancer survival. Nature communications 12, 4098. 10.1038/s41467-021-24343-x.

68. Welsh, R.A., and Sadegh-Nasseri, S. (2020). The love and hate relationship of HLA-DM/DO in the selection of immunodominant epitopes. Current opinion in immunology 64, 117–123. 10.1016/j.coi.2020.05.007.

69. Wengner, A.M., Hopken, U.E., Petrow, P.K., Hartmann, S., Schurigt, U., Brauer, R., and Lipp, M. (2007). CXCR5-and CCR7-dependent lymphoid neogenesis in a murine model of chronic antigen-induced arthritis. Arthritis Rheum 56, 3271–3283. 10.1002/art.22939.

70. Wilbur, C., Yea, C., Licht, C., Irwin, M.S., and Yeh, E.A. (2019). An upfront immunomodulatory therapy protocol for pediatric opsoclonus-myoclonus syndrome. Pediatric blood & cancer 66, e27776. 10.1002/pbc.27776.

71. Zekeridou, A., and Lennon, V.A. (2019). Neurologic Autoimmunity in the Era of Checkpoint Inhibitor Cancer Immunotherapy. Mayo Clin Proc 94, 1865–1878. 10.1016/j.mayocp.2019.02.003.

