## Supplementary figures and images for "Polyclonal lymphoid expansion drives paraneoplastic autoimmunity in neuroblastoma"

### Supplemental Figure 1

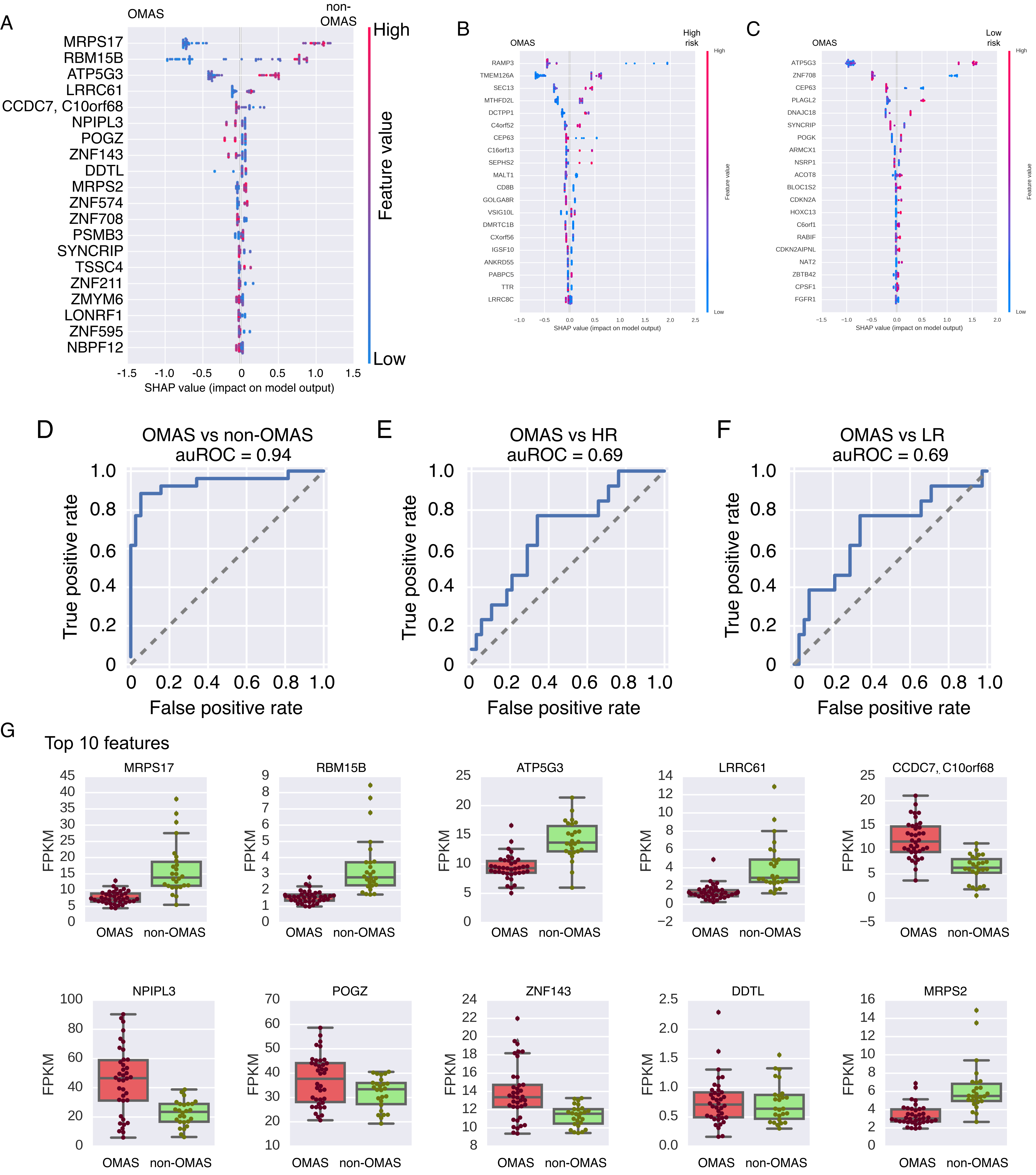


Rosenberg et al. Figure S1

### Supplemental Figure 3

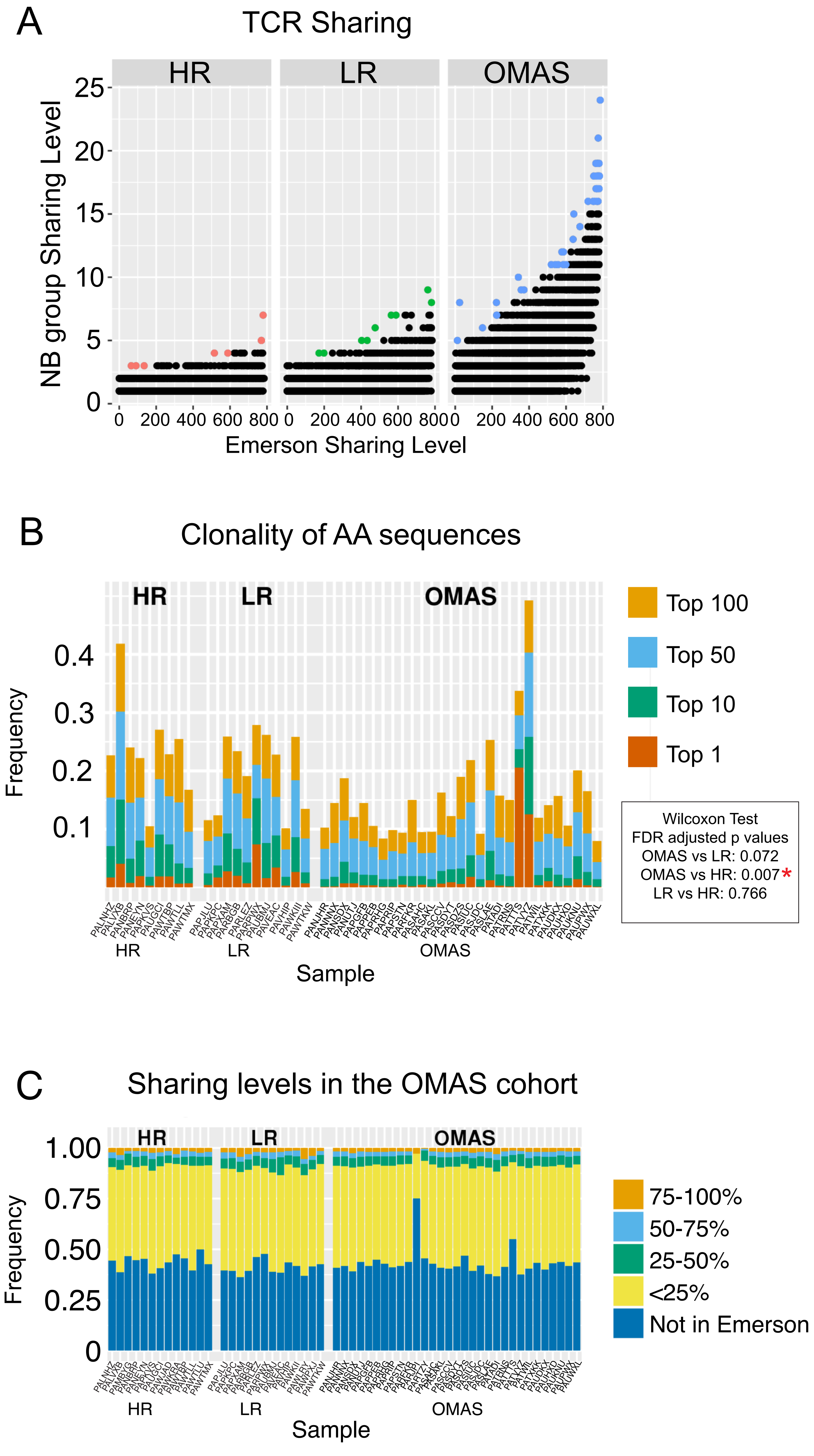


Rosenberg et al Figure S3

### Supplemental Figure 4

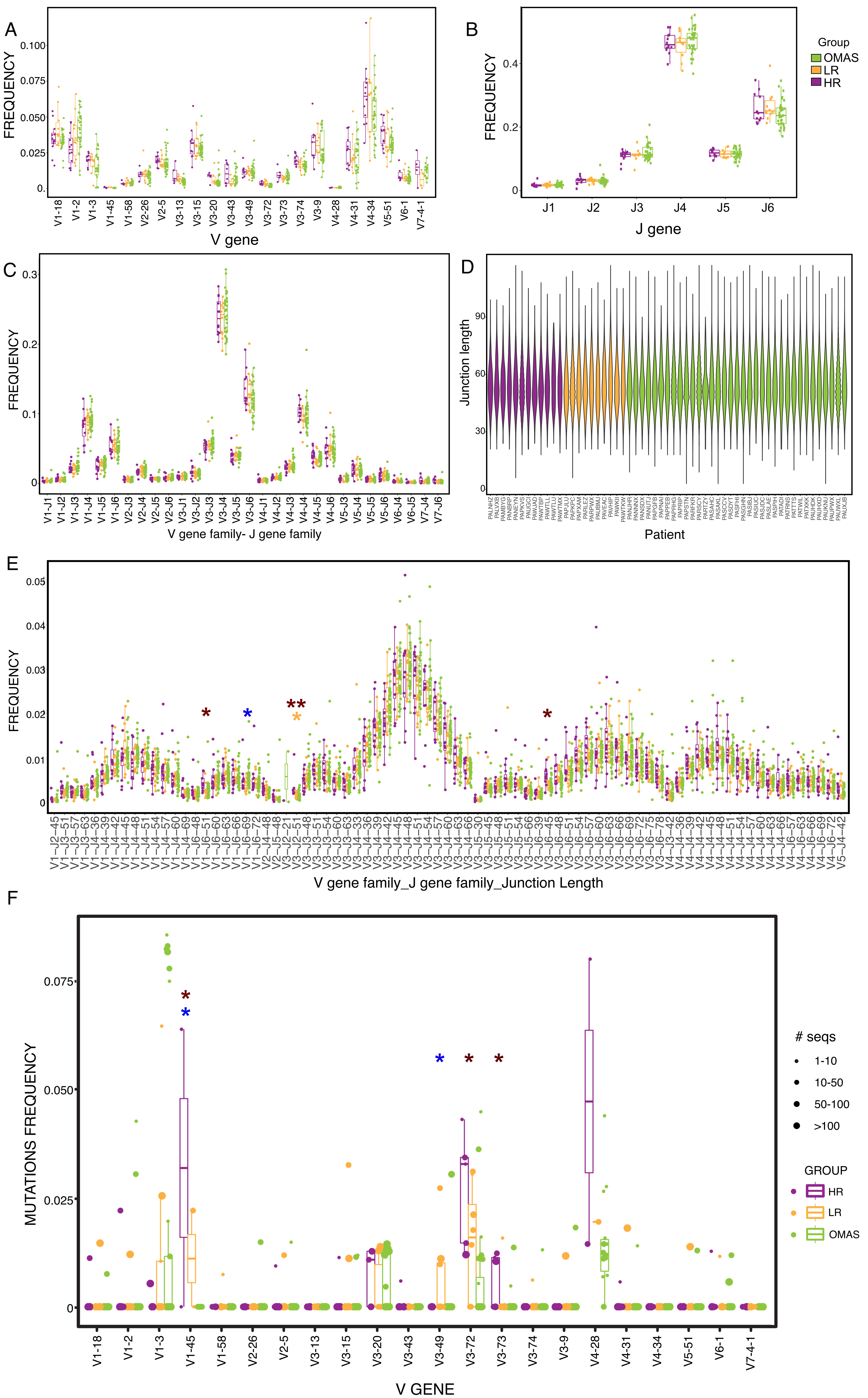


Rosenberg et al Figure S4
