## Supplemental Figure 2 for "Polyclonal lymphoid expansion drives paraneoplastic autoimmunity in neuroblastoma"

A

| Gene ID | Spearman R | Spearman p-value | Gene Symbol | Gene name | Gene function (via genecards) |
|---|---|---|---|---|---|
| 152 | 0.607440624 | 5.25039E-05 | **ADRA2C** | Alpha-2 Adrenergic Receptor Subtype C4 | Alpha-2 adrenergic receptors mediate the catecholamine-induced inhibition of adenylate cyclase through the action of G proteins. These receptors have a critical role in regulating neurotransmitter release from sympathetic nerves and from adrenergic neurons in the central nervous system. |
| 57613 | 0.606877888 | 5.35805E-05 | **KIAA1467** | Protein FAM234B | putative function in hematopoesis/immune system function |
| 3362 | 0.598486518 | 7.21943E-05 | **HTR6** | Serotonin receptor 6 | Rreceptor for 5-hydroxytryptamine (serotonin), a biogenic hormone that functions as a neurotransmitter, a hormone, and a mitogen. The activity of this receptor is mediated by G proteins that stimulate adenylate cyclase. Controls pyramidal neurons migration during corticogenesis, through the regulation of CDK5 activity (By similarity). Is an activator of TOR signaling |
| 23349 | 0.583791451 | 0.000119332 | **KIAA1045** | PHD Finger Protein 24 | unknown |
| 57453 | 0.572185548 | 0.000174528 | **DSCAML1** | DS Cell Adhesion Molecule Like 1 | Cell adhesion molecule that plays a role in neuronal self-avoidance (PubMed:11453658). Promotes repulsion between specific neuronal processes of either the same cell or the same subtype of cells. Promotes both isoneuronal self-avoidance for creating an orderly neurite arborization in retinal rod bipolar cells and heteroneuronal self-avoidance to maintain mosaic spacing between AII amacrine cells (By similarity). Promotes synaptic connectivity via homophilic interactions (By similarity). |
| 1463 | 0.564131769 | 0.00022535 | **NCAN** | Neurocan | May modulate neuronal adhesion and neurite growth during development by binding to neural cell adhesion molecules (NG-CAM and N-CAM). Chondroitin sulfate proteoglycan; binds to hyaluronic acid. |
| 23507 | -0.54981235 | 0.000349386 | **LRRC8B** | Leucine Rich Repeat Containing 8 VRAC Subunit B | Non-essential component of the volume-regulated anion channel (VRAC, also named VSOAC channel), an anion channel required to maintain a constant cell volume in response to extracellular or intracellular osmotic changes |
| 7200 | 0.546202559 | 0.000389021 | **TRH** | Thyrotropin Releasing Hormone | Component of the hypothalamic-pituitary-thyroid axis, it controls the secretion of thyroid-stimulating hormone (TSH) and is involved in thyroid hormone synthesis regulation. Also a modulator of hair growth. |
| 50651 | 0.544077095 | 0.000414195 | **SLC45A1** | Solute Carrier Family 45 Member 1 | Also called Deleted In Neuroblastoma 5 Protein; Proton-associated glucose transporter in the brain. |
| 285596 | 0.541290141 | 0.000449407 | **FAM153A** | Renal Carcinoma Antigen NY-REN-7 | unknown |
| 11105 | 0.53427695 | 0.000550121 | **PRDM7** | PR/SET Domain 7 | Probable histone methyltransferase. |
| 8174 | 0.534035222 | 0.000553926 | **MADCAM1** | Mucosal Vascular Addressin Cell Adhesion Molecule 1 | Cell adhesion leukocyte receptor expressed by mucosal venules, helps to direct lymphocyte traffic into mucosal tissues including the Peyer patches and the intestinal lamina propria. It can bind both integrin alpha-4/beta-7 and L-selectin, regulating both the passage and retention of leukocytes. Isoform 2, lacking the mucin-like domain, may be specialized in supporting integrin alpha-4/beta-7-dependent adhesion strengthening, independent of L-selectin binding. |
| 128506 | 0.531785513 | 0.000590472 | **OCSTAMP** | Osteoclast Stimulatory Transmembrane Protein | Probable cell surface receptor that plays a role in cellular fusion and cell differentiation. Cooperates with DCSTAMP in modulating cell-cell fusion in both osteoclasts and foreign body giant cells (FBGCs). Involved in osteoclast bone resorption. Promotes osteoclast differentiation and may play a role in the multinucleated osteoclast maturation (By similarity). |
| 339260 | 0.530537824 | 0.000611653 | **uncharacterized** | | ncRNA; uncharacterized |
| 29944 | 0.529796946 | 0.000624548 | **PNMA3** | Paraneoplastic Cancer-Testis-Brain Antigen | The protein encoded by this gene belongs to the paraneoplastic antigen MA (PNMA) family, which shares homology with retroviral Gag proteins. The PNMA antigens are highly expressed in the brain and also in a range of tumors associated with serious neurological phenotypes. |
| 6543 | 0.525245675 | 0.00070921 | **SLC8A2** | Solute Carrier Family 8 Member A2 | Mediates the electrogenic exchange of Ca(2+) against Na(+) ions across the cell membrane, and thereby contributes to the regulation of cytoplasmic Ca(2+) levels and Ca(2+)-dependent cellular processes. Contributes to cellular Ca(2+) homeostasis in excitable cells. Contributes to the rapid decrease of cytoplasmic Ca(2+) levels back to baseline after neuronal activation, and thereby contributes to modulate synaptic plasticity, learning and memory. Plays a role in regulating urinary Ca(2+) and Na(+) excretion. |
| 644150 | 0.521164598 | 0.000793648 | **WIPF3** | WAS/WASL Interacting Protein Family Member | May be a regulator of cytoskeletal organization. May have a role in spermatogenesis (By similarity). |
| 54020 | 0.52008652 | 0.000817394 | **SLC37A1** | Glycerol-3-Phosphate Permease | Inorganic phosphate and glucose-6-phosphate antiporter. May transport cytoplasmic glucose-6-phosphate into the lumen of the endoplasmic reticulum and translocate inorganic phosphate into the opposite direction. Independent of a lumenal glucose-6-phosphatase. May not play a role in homeostatic regulation of blood glucose levels. |
| 2645 | 0.519269593 | 0.000835806 | **GCK** | Glucokinase | Catalyzes the phosphorylation of hexose, such as D-glucose, D-fructose and D-mannose, to hexose 6-phosphate (D-glucose 6-phosphate, D-fructose 6-phosphate and D-mannose 6-phosphate, respectively) |
| 164395 | 0.518878361 | 0.000844753 | **TTLL9** | Tubulin Tyrosine Ligase Like 9 | Probable tubulin polyglutamylase that forms polyglutamate side chains on tubulin. |
| 219770 | 0.518326773 | 0.000857511 | **GJD4** | Gap Junction Protein Delta 4 | One gap junction consists of a cluster of closely packed pairs of transmembrane channels, the connexons, through which materials of low MW diffuse from one cell to a neighboring cell. |
| 3776 | -0.51813137 | 0.000862072 | **KCNK2** | K2P2.1 Potassium Channel | Ion channel that contributes to passive transmembrane potassium transport (PubMed:23169818). In astrocytes, the heterodimer formed by KCNK1 and KCNK2 is required for rapid glutamate release in response to activation of G-protein coupled receptors, such as F2R and CNR1 (By similarity). |
| 55301 | -0.51809723 | 0.000862871 | **OLAH** | Oleoyl-ACP Hydrolase | Contributes to the release of free fatty acids from fatty acid synthase (FASN). Has broad substrate specificity, giving rise to a range of free fatty acids with chain lengths between 10 and 16 carbon atoms (C10 - C16). |
| 3906 | 0.509851101 | 0.001076247 | **LALBA** | Lactalbumin Alpha | Regulatory subunit of lactose synthase, changes the substrate specificity of galactosyltransferase in the mammary gland making glucose a good acceptor substrate for this enzyme. This enables LS to synthesize lactose, the major carbohydrate component of milk. In other tissues, galactosyltransferase transfers galactose onto the N-acetylglucosamine of the oligosaccharide chains in glycoproteins. |
| 154790 | 0.508510102 | 0.001115043 | **CLEC2L** | C-Type Lectin Domain Family 2 Member L | putative carbohydrate binding |
| 23217 | 0.50588836 | 0.001194483 | **ZFR2** | Zinc Finger RNA Binding Protein 2 | putative RNA binding |
| 26240 | 0.501586708 | 0.001335698 | **FAM50B** | XAP5-Like Protein | Imprinted. Promoter methylation of the maternal allele may restrict expression to the paternal allele in placenta. |

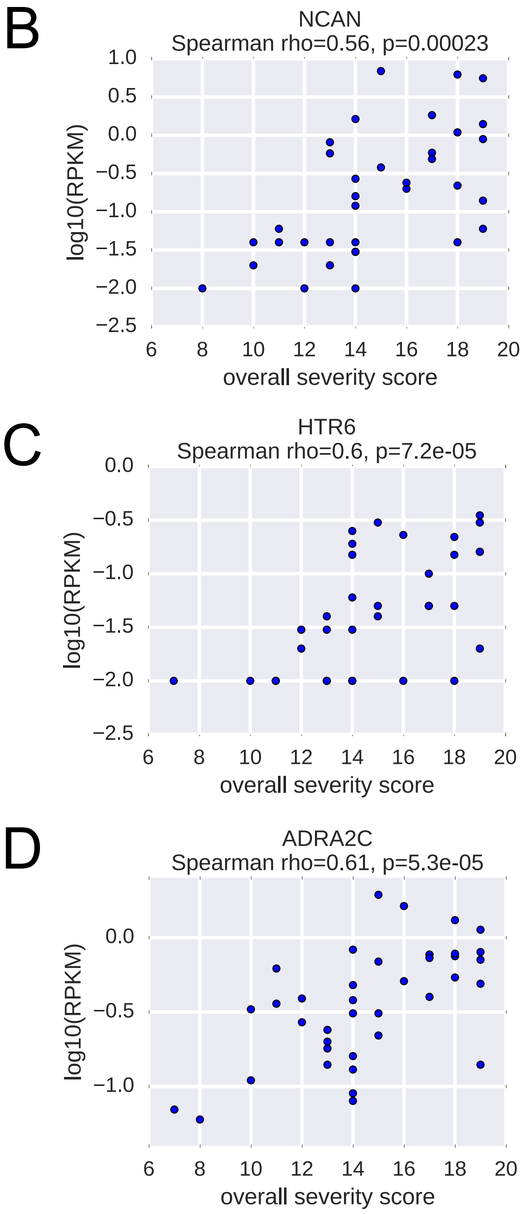


Rosenberg et al Figure S2
