## Supplemental Figure 5 for "Polyclonal lymphoid expansion drives paraneoplastic autoimmunity in neuroblastoma"

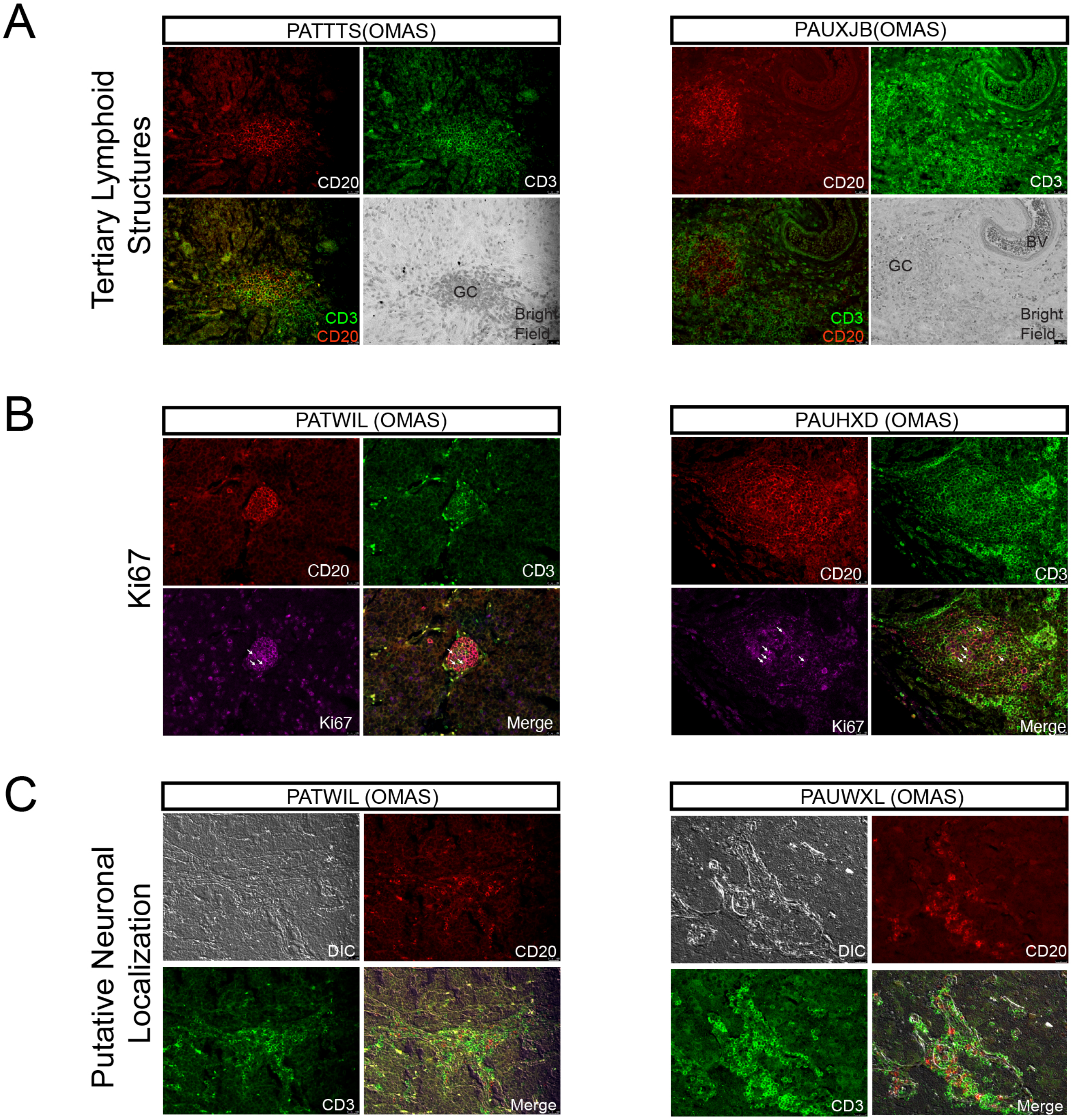


D

| patient USI | Tumor type | Follicles | Germinal centers | Dominant CD20 vs CD3 | Neuropile | Homer-Wright rosettes | Diff. ganglilar neuronal cells? | comments for column H | additional comments | patient group |
|---|---|---|---|---|---|---|---|---|---|---|
| PANBRP | NB | 0 | 0 | none | yes | yes | no | | | HR |
| PAUGCI | NB | 0 | 0 | none | no | no | no | | MYCN amplified | HR |
| PAWTMX | NB | 0 | 0 | none | yes | no | yes | | | HR |
| PAPKVS | GNB | 1+ | 1+ | CD20 | yes | no | yes | dominant cell type | MYCN amplified | HR |
| PANEYN | NB | 3+ | 3+ | CD20 | no | yes | yes | but <50% tumor | | HR |
| PAPKPC | GNB | 0 | 0 | none | yes | yes | yes | dominant cell type | | LR |
| PAPXAM | GNB | 0 | 0 | none | no | no | no | | | LR |
| PAVHIP | NB | 1+ | 0 | ?very weak | yes | no | no | | | LR |
| PARPWX | NB | 1+ | 1+ | CD20 | yes | no | no | | | LR |
| PAVEAC | GNB | 2+ | 2+*** | CD20 weak | yes | no | yes | dominant cell type | ***: F&GC in connection with NB component, as also seen in other GNBs | LR |
| PAPJLU | GNB | 3+ | 3+ | CD20 | yes | yes | yes | | | LR |
| PAPRIP | GNB | 0 | 0 | none | yes | no | yes | only few in the slide though* | *: probably dominant in other slides from tumor since it's GNB | OMAS |
| PATNJJ | NB | 0 | 0 | none | yes | no | no | | | OMAS |
| PANUTJ | NB | 0 | 0 | none | no | yes | no | | | OMAS |
| PASDYT | NB | 1+ | 1+ | CD20 | yes | no | no | | | OMAS |
| PASAKL | NB | 2+ | 2+ | CD20 | yes | no | yes | just very few | | OMAS |
| PASJDC | NB | 2+ | 2+ | CD20 (+CD8?) | yes | no | no | | | OMAS |
| PATTTS | GNB | 2+ | 2+ | CD20 | yes | no | yes | dominant cell type | | OMAS |
| PATWIL | NB | 2+ | 2+ | CD20 | no | yes | no | | | OMAS |
| PANJHR | NB | 3+ | 3+ | CD20 | no | no | yes | | | OMAS |
| PAPPEB | NB | 3+ | 3+ | CD20 | yes | no | yes | | | OMAS |
| PAPSTN | NB | 3+ | 3+ | CD20+CD3 | no | no | no | | | OMAS |
| PAUHXD | NB | 3+ | 3+ | CD20 | no | yes | no | | | OMAS |
| PAUWXL | NB | 3+ | 3+ | CD20 | yes | no | yes | very few | | OMAS |
| PAUXJB | GNB | 3+ | 3+ | CD20** | yes | no | yes | | **: Many interfollicular CD3+ T lymphocytes | OMAS |

Rosenberg et al Figure S5
